# Neuronal genetic rescue normalizes brain network dynamics in a lysosomal storage disorder despite persistent storage accumulation

**DOI:** 10.1101/2021.05.03.442437

**Authors:** Rebecca C. Ahrens-Nicklas, Luis Tecedor, Arron F. Hall, Owen Kane, Richard J. Chung, Elena Lysenko, Eric D. Marsh, Colleen S. Stein, Beverly L. Davidson

## Abstract

Although neurologic symptoms occur in two-thirds of lysosomal storage disorders (LSDs), for most we do not understand the mechanisms underlying brain dysfunction. A major unanswered question is if the pathogenic hallmark of LSDs, storage accumulation, induces functional defects directly or is a disease bystander. Also, for most LSDs we do not know the impact of loss-of-function in individual cell types. Understanding these critical questions are essential to therapy development. Here, we determined the impact of genetic rescue in distinct cell types on neural circuit dysfunction in CLN3 disease, the most common pediatric dementia and a representative LSD. We restored *Cln3* expression via AAV-mediated gene delivery and conditional genetic rescue in a CLN3 mouse model. Surprisingly, we found that low-level rescue of *Cln3* expression in neurons alone normalized clinically-relevant electrophysiologic markers of network dysfunction, despite the presence of substantial residual histopathology, in contrast to restoring expression in astrocytes. Thus, loss of Cln3 function in neurons, not storage accumulation, underlies neurologic dysfunction in CLN3 disease, implying that storage clearance may be an inappropriate target for therapy development and an ineffectual biomarker.

## Introduction

Lysosomal storage disorders (LSDs) are a group of approximately 50 rare metabolic diseases characterized by progressive accumulation of storage material in lysosomes. Neurological symptoms, including neurocognitive regression, seizures, and psychiatric symptoms, occur in more than 2/3 of LSDs and are generally not ameliorated by current therapies ^1^. For many LSDs, including CLN3 disease, the most common cause of childhood dementia, the mechanisms underlying neurologic dysfunction are poorly understood.

The function of CLN3 protein remains unknown, hindering mechanism-based therapy development. The brains of CLN3 disease mouse models demonstrate progressive storage accumulation, predominantly comprised of subunit C of the mitochondrial ATP synthase (SCMAS) ^2 3^, mirroring what occurs in CLN3 disease patients^4^. Therefore, most drug development efforts rely on histopathologic markers of efficacy, including reduction of storage material, reactive astrocytosis and microgliosis^2,4,5^. However, whether these histopathologic changes cause disease, augment the disease process, or are simply harmless epiphenomenon is unclear.

In addition to lack of clarity on the impact of storage, the role of various CNS cell types in driving functional deficits in CLN3 disease is unknown. Astrocytes and microglia isolated from CLN3 disease mouse models demonstrate altered cellular properties and physiology^6-8^, and in animal models, localized glial activation occurs in regions where neurons will later die^9^ supporting a role for glia as a disease driver. On the other hand, neurons from CLN3 animal models exhibit both histopathologic and functional differences^2,10,11^. Ultimately, patients’ symptoms such as seizures and neurologic regression arise directly from functional changes in key neuronal circuits. Nonetheless, we do not understand what cell types underlie these pathologic network dynamics in the brain.

Previously, we described network-level neurologic defects in both the cortex and hippocampus of two CLN3 disease mouse models using *in vitro* voltage sensitive dye imaging and *in vivo* electrophysiology techniques^12^. While many brain regions are affected in CLN3 disease, the cortex and hippocampus are especially vulnerable and show early pathologic changes in mouse models and human patients^3,4,11,13^. We found that network-level changes begin early in the disease course before substantial storage accumulation in both the hippocampus and cortex. In slice-recordings, these changes include hypoexcitability of the dentate gyrus of the hippocampus, a region essential for learning and memory. On *in vivo* electroencephalogram (EEG), CLN3 disease mice show frequent epileptiform spikes, a feature of many epilepsy models, and a shift towards high frequency background activity, a change associated with cognitive impairment and present in Alzheimer’s disease models and patients^14,15^.

To date, CLN3 disease research has been hindered by the fact that animal models have only subtle behavioral and survival phenotypes^16,17^. Most manifest histopathologic changes including storage accumulation and reactive astrocytosis; however, these phenotypes may not correlate with central nervous system (CNS) dysfunction. Our network-level electrophysiologic approach provides a robust, clinically meaningful measure of neurologic function that can probe disease mechanisms and evaluate potential therapies.

Here, we developed a new ‘conditional rescue’ mouse model of CLN3 disease. The model is homozygous for the most common human CLN3 disease mutation^2^, but also carries a Cre-inducible wild-type *Cln3* transgene, under the control of its endogenous promoter, allowing for cell-type specific rescue of expression. We employed AAV-mediated gene replacement as a second strategy to restore *Cln3* expression and combined these tools with network-level electrophysiology techniques to answer three essential questions: 1) Does gene replacement correct neural circuit dysfunction in a LSD? 2) What cell types must be corrected to improve network dynamics? and 3) Does storage accumulation underlie deficits in CNS network dynamics in CLN3 disease?

## Results

In prior work, we found that CLN3 disease mouse models demonstrate robust, progressive neuronal network defects^12^. We first assessed if neonatal correction of most CNS cell types could prevent circuit-level functional pathology. For this, we employed an adeno-associated virus serotype 9 (AAV9) vector expressing wild-type murine *Cln3* under control of 1665 bp of the endogenous *Cln3* promoter, which mimics the low levels of expression seen in the endogenous setting **(Sup Figure 1-2, Sup Methods)**. Vector was delivered at p0 via unilateral intracerebroventricular (ICV) injection into CLN3-deficient mice to transduce both astrocytes and neurons^18^ **(Sup Figure 3)**. The CLN3 disease mouse model harbors the most common human CLN3 disease mutation, a deletion of exons 7 and 8 (*Cln3*^Δex78/Δex78^) (*2*). ICV injection of AAV9.CLN3 at a dose of 1e^10^ vg increased *Cln3* expression **(Figure 1A)**. One year after injection, levels in the *Cln3*^Δex78/Δex78^ hemisphere contralateral to the injection site reached 24.5% ± 4.7 of heterozygous controls. When assessed histologically, gene replacement partially reduced storage accumulation in both the hippocampus and motor cortex ipsilateral to the injection site **(Figure 1B, Sup Figure 4**).

**Figure 1:**
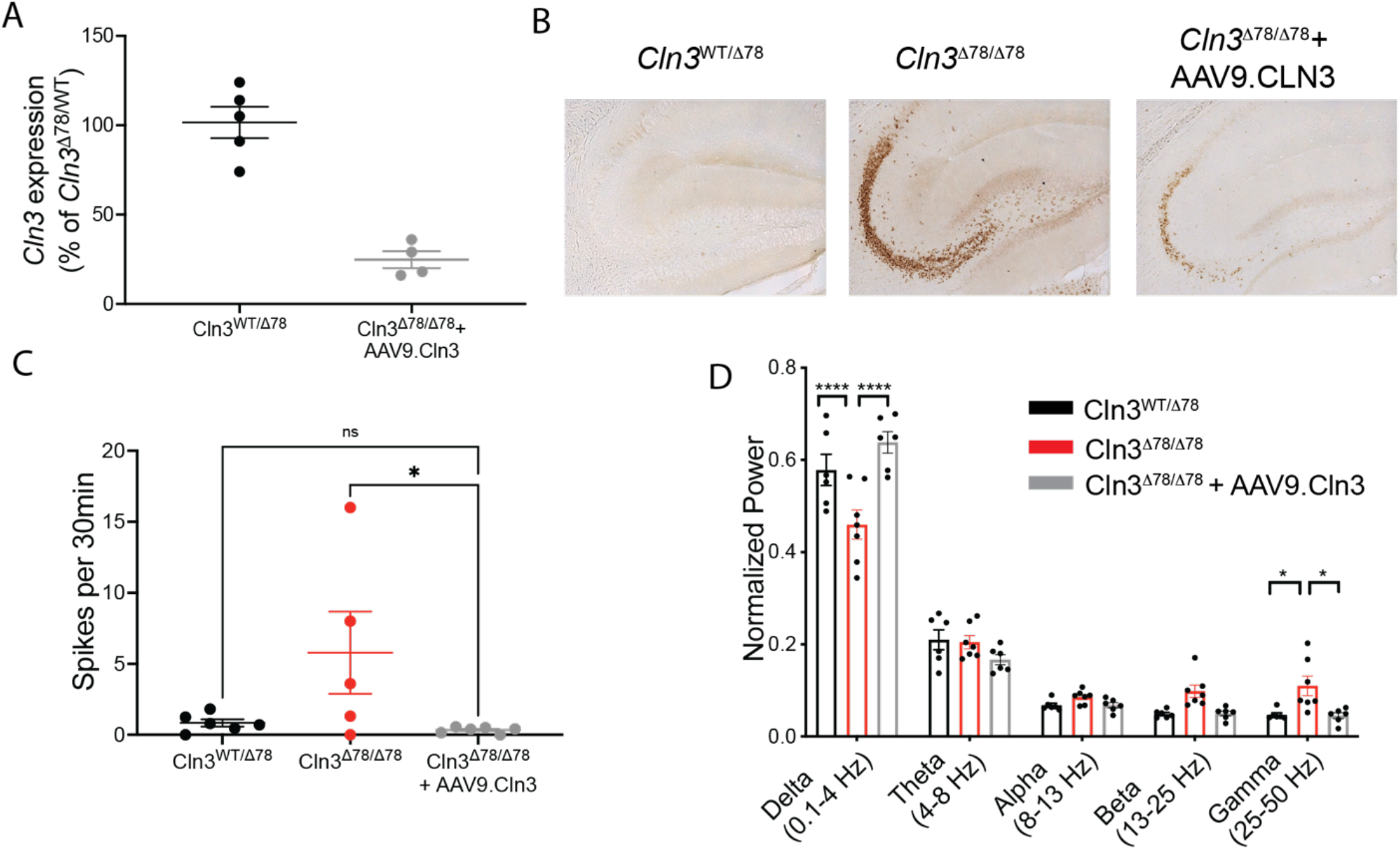
Intracerebroventricular AAV9.CLN3 gene therapy given at p0 normalizes neuronal network dynamics *in vivo* at 12 months of age. **(A)** p0 ICV injection of AAV9.Cln3 restored Cln3 expression in *Cln3*^Δex78/Δex78^ mouse brain at 12 months of age, with levels in the *Cln3*^Δex78/Δex78^ hemisphere contralateral to the injection site reaching 24.5% ± 4.7 of heterozygous controls. **(B)** Subunit C of the mitochondrial ATP synthase (SCMAS) accumulation is partially corrected in the hippocampus ipsilateral to the injection site of *Cln3*^Δex78/Δex78^ mice treated with AAV9.CLN3. Gene replacement corrects circuit function measured on intracranial EEG including **(C)** epileptiform spiking rates throughout the cortex (average of all cortical leads shown) and **(D)** background frequency band composition in the motor cortex. Groups: *Cln3*^WT/Δex78^ (black); *Cln3*^Δex78/Δex78^ (red); *Cln3*^Δex78 /Δex78^ + AAV9.CLN3 (gray). N=5-6 animals/condition. Results from 12 hours of recordings were averaged for each animal shown as an individual circle. Spikes analyzed using a non-parametric Kruskal-Wallis test followed by Dunn’s multiple comparisons test. Frequency bands were analyzed by two-way ANOVA by band and genotype, followed by Bonferonni’s multiple comparisons test (significant results of multiple comparison testing shown as \**p*<0.05, \*\**p*<0.01, \*\*\**p*<0.001,\*\*\*\**p<*0.0001). Data are shown as mean +/-s.e.m.

To determine the impact of p0 AAV9.CLN3 infusion on *in vivo* network dynamics, we recorded activity throughout the brain via *in vivo* EEG. Twelve-month-old *Cln3*^Δex78/Δex78^ mice have increased spike burden and a shift in background activity towards faster frequencies, with increased power in faster frequency bands, as compared to *Cln3*^Δex78/WT^ controls **(Figure 1C-D)**. Specifically, there is decreased power in the slow delta frequency band (0.1-4Hz) and a trend towards increased power in the faster beta (13-25Hz) and gamma (25-50Hz) bands in *Cln3*^Δex78/Δex78^ mice, consistent with our previous report in *Cln3* knock-out mice^12^. Remarkably, restoring gene expression at p0 normalized spiking rates **(Figure 1C)** and corrected background activity **(Figure 1D)** in 12-month-old *Cln3*^Δex78/Δex78^ AAV9.CLN3-treated animals. Correction of EEG phenotypes was equally effective in both the hemispheres. Thus, early gene replacement can normalize network dynamics, and this rescue does not require full inhibition of storage accumulation.

Because AAV9 delivered to neonates transduces astrocytes and neurons^18^, it is unclear whether all cell types must be corrected to improve circuit function, a question critically important to translate gene therapies to children. AAV9 delivered into the CNS generally transduces neurons in adult mice and juvenile and adult large animal models^19-21^. To address this, we developed a conditional rescue model of CLN3 disease, the Flex*Cln3* mouse. This model is homozygous for the common *Cln3* exon 7-8 deletion and also carries the Cre-inducible Flex*Cln3* allele, permitting expression of wild-type Cln3 protein in the presence of Cre recombinase **(Figure 2, Sup Methods and Sup Figures 5-7)**. The Flex*Cln3* allele is under control of the same 1665 bp *Cln3* promoter as used for AAV-mediated gene replacement.

**Figure 2:**
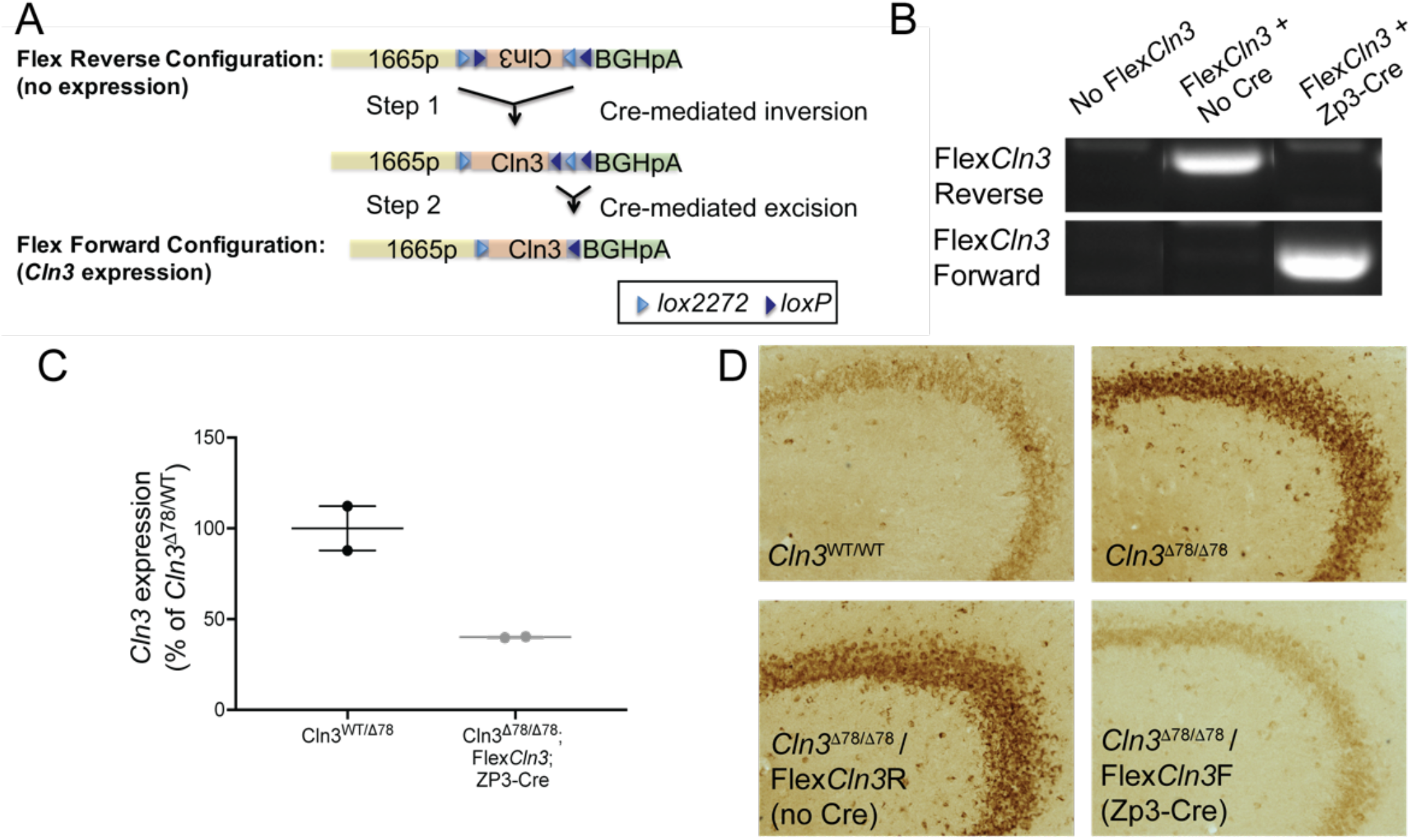
Expression of Flex*Cln3* allele in all cells reduces storage accumulation in *Cln3*^Δex78/Δex78^ brain. **(A)** In the presence of Cre, the Flex*Cln3* allele undergoes inversion (Step 1) and excision (Step 2) and is expressed under control of its endogenous promoter. **(B)** PCR of genomic DNA from hippocampus using orientation-specific primers confirms that the Flex*Cln3* is in the forward orientation only in the presence of Cre (see Sup Figure 7 for uncut gels). **(C)** In the presence of *Zp3*-Cre which drives ubiquitous recombination, the Flex*Cln3* allele results in expression levels that are 40.0 +/-0.4% of heterozygous controls. **(D)** Subunit C of the mitochondrial ATP synthase (SCMAS) accumulates in the hippocampus of *Cln3*^Δex78/Δex78^ and Flex*Cln3* / *Cln3*^Δex78/Δex78^ mice. Storage is reduced in mice with the Flex*Cln3* allele in the forward confirmation.

We validated the model by crossing Flex*Cln3* mice with *Zp3-Cre* mice^22^ to drive germline level recombination and flip the reverse-oriented wildtype Flex*Cln3* allele into the forward orientation in all cells. Recombination of the allele was confirmed using orientation-specific PCR primers on genomic DNA extracted from the hippocampus **(Figure 2B, Sup Figure 7)**. In these mice *Cln3* expression was ubiquitously restored, with expression levels dictated by the transgenic *Cln3* 1665 bp promoter. *Cln3* expression in the hippocampus was restored to 40% of heterozygous (*Cln3*^Δex78/WT^) control levels as determined by qPCR **(Figure 2C)**. Correction (i.e. expression of the Flex*Cln3* transgene) in all cell types (Flex*Cln3*/ *Cln3*^Δex78/Δex78^/ *Zp3-*Cre) substantially reduced SCMAS storage in the hippocampus and other brain regions **(Figure 2D, Sup Figure 8)**.

For selective rescue of astrocytes or neurons, Flex*Cln3* mice were crossed to *Gfap-*Cre^23^ or *Syn1-*Cre^24^ driver lines, respectively. Cre-mediated rearrangement of the Flex*Cln3* allele was confirmed by PCR of genomic DNA extracted from the hippocampus using orientation specific primers **(Figure 3A)**. As Cre is only expressed in a subset of cells (either astrocytes or neurons), unlike in the case of *Zp3*-Cre **(Figure 2B)**, both the forward and reverse alleles were detected in hippocampal tissue.

**Figure 3:**
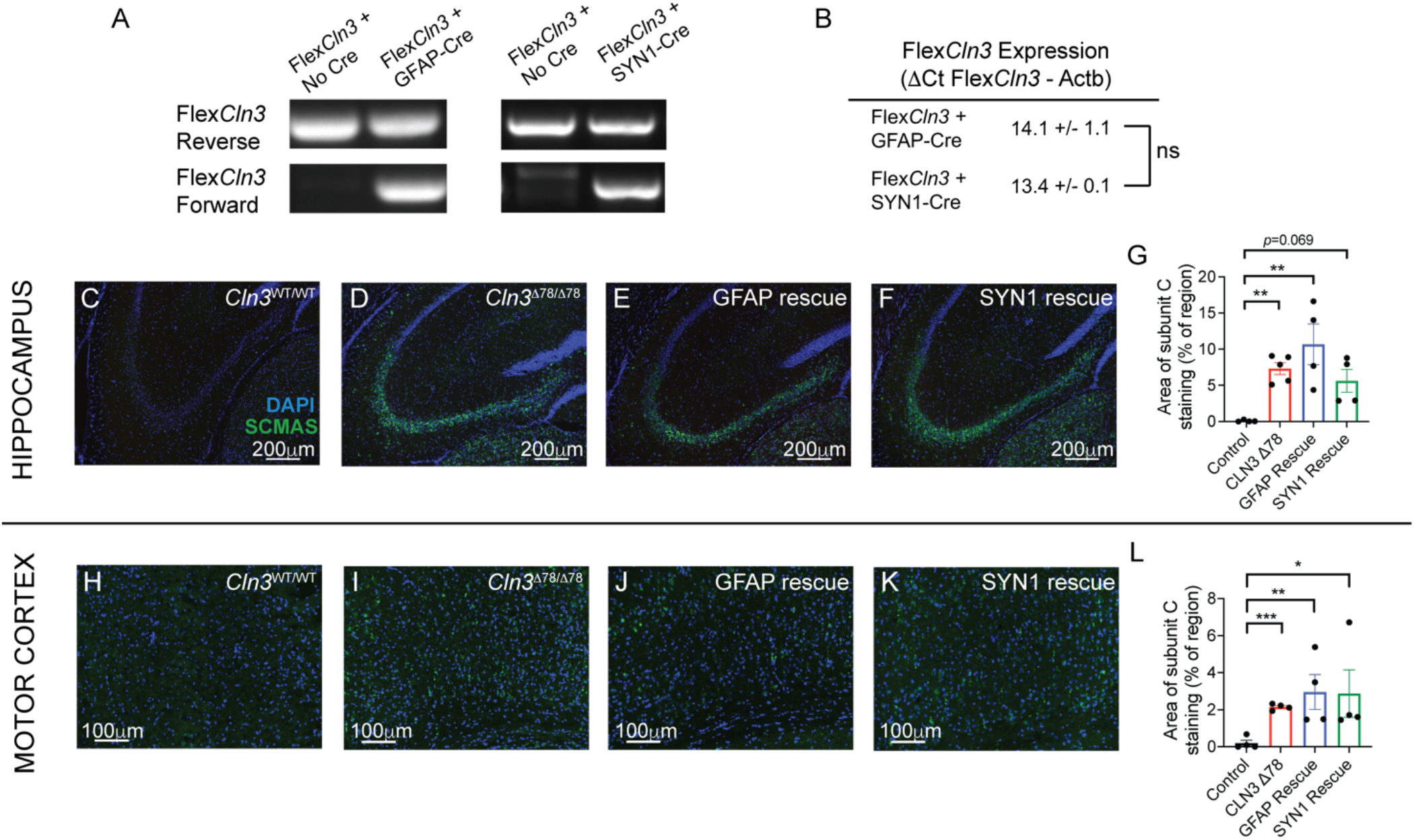
Expression of Flex*Cln3* allele in either neurons or astrocytes alone does not prevent storage accumulation. **(A)** In the presence of Cre, the Flex*Cln3* allele undergoes inversion as detected by PCR with orientation-specific primers of genomic DNA from hippocampus. As both the *Gfap*-Cre and *Syn1*-Cre only drive recombination in a subset of cells, both the forward and reverse alleles are detected (see Sup Figure 7 for uncut gels). **(B)** Expression levels as measured by ΔCt values (FlexCln3-Δ actin) are similar in Flex*Cln3* brain in the presence of *Gfap*-Cre and *Syn1*-Cre (14.1 vs. 13.4, *p*=0.53 by unpaired t-test). **(C-G)** Expression of Flex*Cln3* in astrocytes (Flex*Cln3* / *Cln3*^Δex78/Δex78^ / *Gfap-*Cre animals) or neurons alone (Flex*Cln3* / *Cln3*^Δex78/Δex78^ / *Syn1-*Cre animals) does not prevent accumulation of SCMAS (green). **(H-K)** Similar findings were seen in the motor cortex. N=4-5 animals, non-parametric Kruskal-Wallis test followed by Benjamini and Hochberg multiple comparisons correction to detect differences from control. For all panels: \**p*<0.05, \*\**p*<0.01, \*\*\**p*<0.001, \*\*\*\**p*<0.0001.

Expression of forward-oriented Flex*Cln3* transcripts was confirmed in the hippocampus of the astrocyte and neuronal-specific rescue models. As rescue did not occur in all cell types, and Cln3 protomer activity and expression can vary between cells and be induced by disease pathology^3^ it is difficult to normalize expression levels to heterozygote controls. However, to get a relative sense of expression levels we compared the level of Flex*Cln3* transgene expression in the astrocyte-only and neuron-only rescue models. We found no differences in transcript levels (normalized ΔCt values of 14.1 vs. 13.4, *p*=0.53 by unpaired t-test) between the groups **(Figure 3B)**. As there are no available antibodies that can reliably detect endogenous levels of mouse Cln3 protein^25^, we could not assess protein expression.

We next determined the impact of cell type-selective restoration of *Cln3* on histopathological readouts. Rescue of *Cln3* expression in either astrocytes (Flex*Cln3* / *Cln3*^Δex78/Δex78^ / *Gfap-*Cre) or neurons (Flex*Cln3* / *Cln3*^Δex78/Δex78^ / *Syn1-*Cre) alone did not prevent storage accumulation in the hippocampus or motor cortex in 6 or 12-month-old mice **(Figure 3C-L, Sup Figures 8-9)**. In addition to storage accumulation, CLN3 disease mice develop progressive, reactive astrocytosis. At twelve months of age, both the hippocampus and motor cortex of *Cln3*^Δex78/Δex78^ showed increased astrocytosis as compared to wild-type controls. Rescue of *Cln3* expression in either neurons or astrocytes alone did not prevent astrocytosis **(Sup Figure 10)**.

To investigate if the Flex*Cln3* transgene could prevent storage accumulation in the specific cell type where it was expressed, we investigated co-localization of autofluorescent lipofuscin with a neuronal marker, NeuN, in cortical tissue **(Figure 4)**. There is little lipofuscin accumulation in *Cln3*^WT/WT^ cortex **(Figure 4, top row)**, as compared to *Cln3*^Δex78/Δex78^ cortex, where there is storage in both neurons and non-neuronal cells (**Figure 4, second row)**. Ubiquitous Flex*Cln3* allele expression (in the presence of *Zp3*-Cre) prevented storage accumulation in all cell types **(Figure 4, third row)**. Flex*Cln3* allele expression in neurons (in the presence of *Syn1*-Cre) did not prevent storage in either neurons or non-neuronal cells **(Figure 4, bottom row)**.

**Figure 4:**
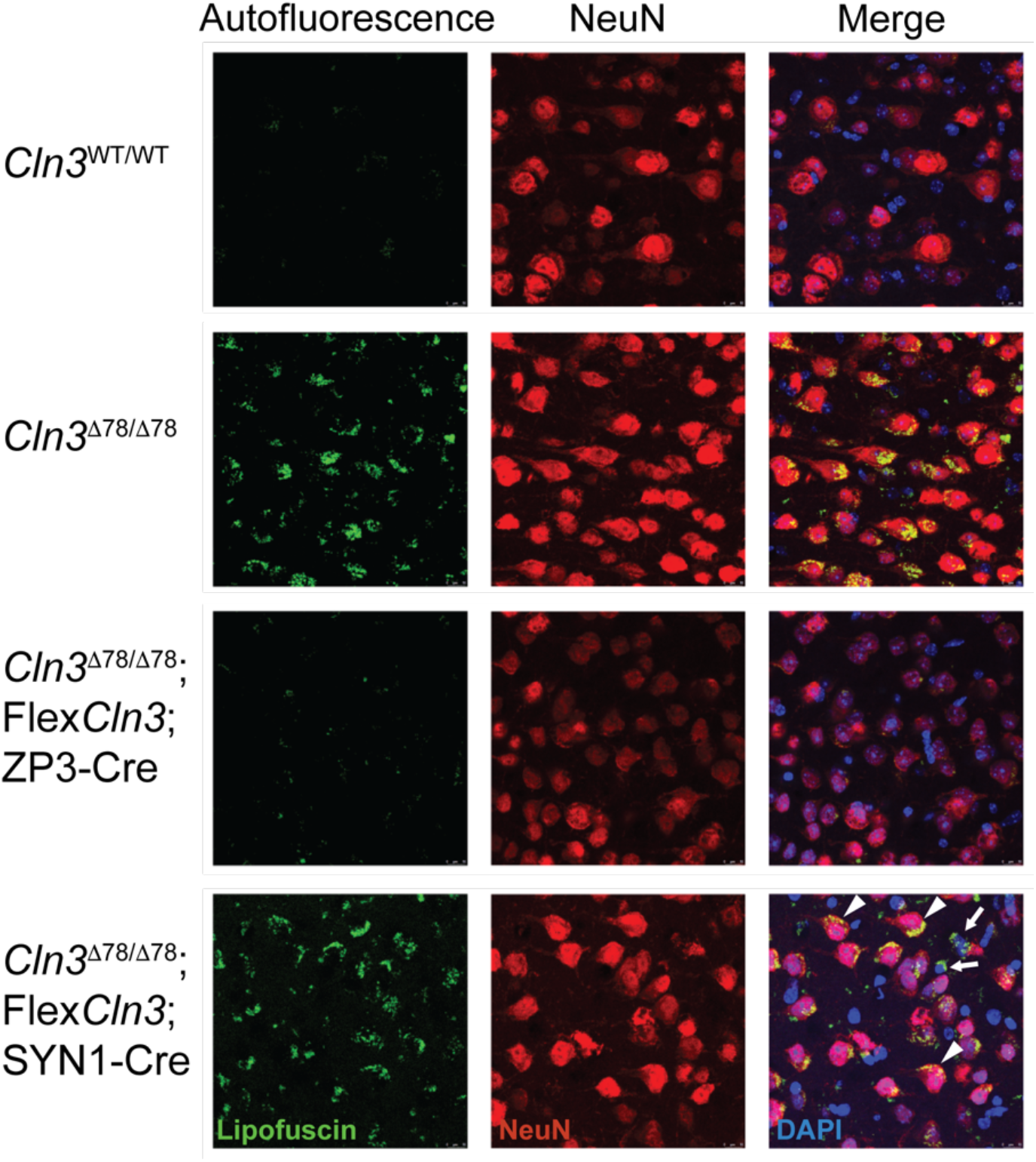
Expression of the Flex*Cln3* allele in neurons alone does not prevent autofluorescent lipofuscin accumulation in cortical neurons. **(Top row)** There is little autofluorescent lipofuscin accumulation (green) in wild-type cortical neurons (red) (blue DAPI) as compared to **(second row)** *Cln3*^Δ 78/Δ 78^ cortical neurons in 12 month old animals. **(Third row)** *Zp3*-Cre induced expression of the Flex*Cln3* allele in all cell types prevents autofluorescent lipofuscin accumulation in both neuronal and non-neuronal cells. **(Bottom row)** Flex*Cln3* expression in neurons (*Syn1*-Cre) did not prevent storage accumulation in neurons (arrow heads) or non-neuronal cells (arrows).

Next, we determined if restoring *Cln3* expression could improve electrophysiologic measures, despite residual storage accumulation. *In vitro* voltage sensitive dye imaging of hippocampal slices from 6-month-old animals was completed in Flex*Cln3* / *Cln3*^Δex78/Δex78^ / *Gfap-*Cre, Flex*Cln3* / *Cln3*^Δex78/Δex78^ / *Syn1-*Cre, *and* Flex*Cln/Cln3*^Δex78/Δex78^ /Zp3 mice. In this early-stage of disease, the internal blade of the dentate gyrus of the *Cln3*^Δex78/Δex78^ hippocampus was hypoexcitable in response to perforant path stimulation as compared to heterozygous *Cln3*^Δ 78/WT^ mice **(Figure 5A-C)**, consistent with our previous studies^12^. Specifically, 32.7% of pixels in the internal blade of the *Cln3*^Δex78/Δex78^ dentate gyrus were significantly hypoexcitable as compared to slices obtained from heterozygous control animals. Interestingly, restoring expression in astrocytes did not correct hypoexcitability of the dentate gyrus **(Figure 5D-E)**. However, restoring *Cln3* expression in neurons was sufficient to prevent disease-induced hypoexcitability **(Figure 5F-G)**, in fact excitability was mildly increased as compared to controls. This rescue of dynamics occurred despite the presence of residual histopathology **(Figure 3-4 and Sup. Figures 8-10)**.

**Figure 5:**
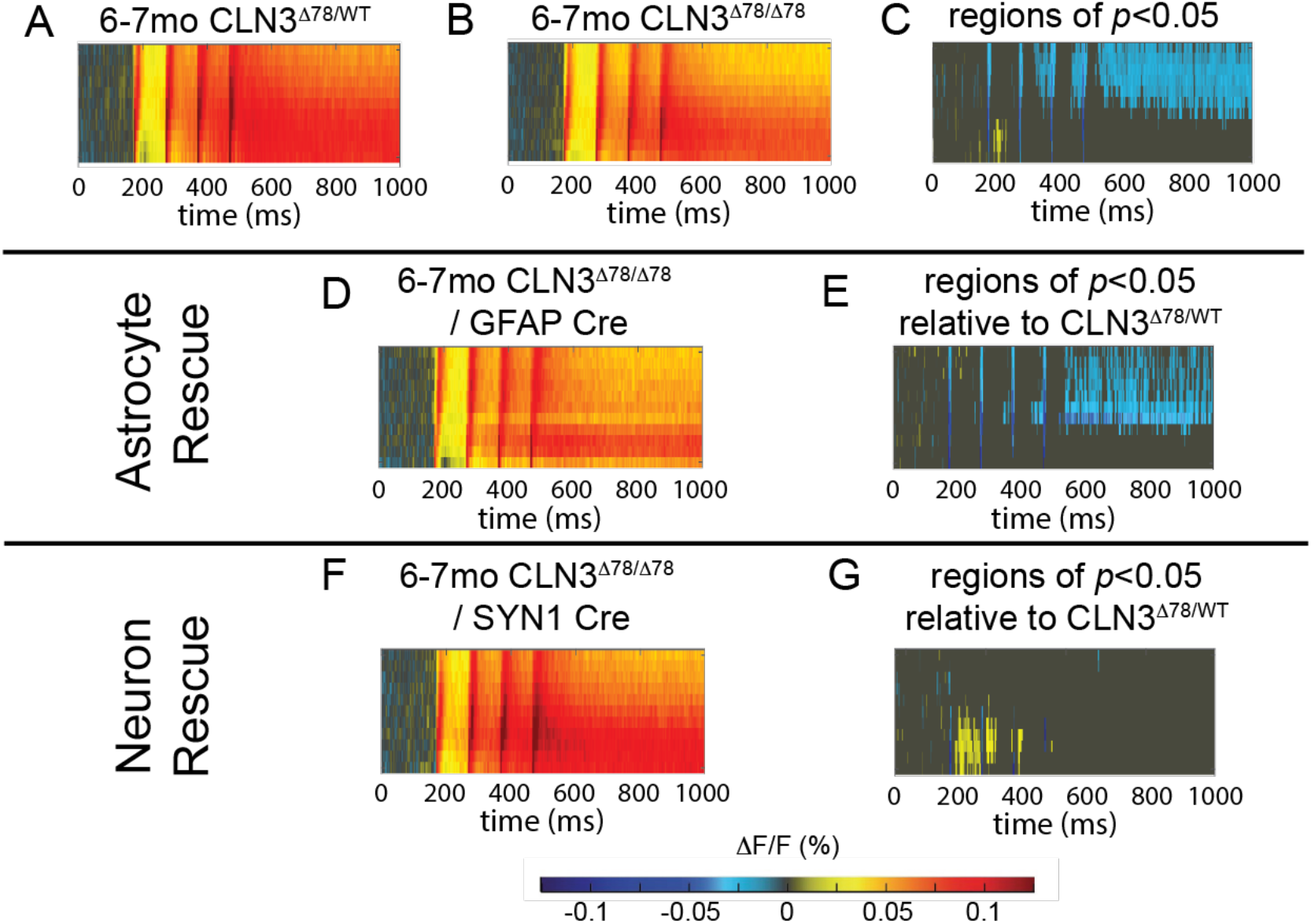
Restoring low level *Cln3* expression in neurons alone corrects hypoexcitability of the dentate gyrus. Dentate gyrus excitability measured through *in vitro* voltage sensitive dye imaging of hippocampal slices from 6-month-old animals is shown as raster plots of average fluorescence change (Δ F/F warm colors excitation, cool colors inhibition) over time (X axis) and location within the internal blade of the dentate gyrus (Y Axis). **(A)** Four-pulse stimulation of the perforant path in heterozygous control (*Cln3*^Δex78/WT^) mice reveals robust excitation of the dentate gyrus (DG) as compared to **(B)** *Cln3*^Δex78/Δex78^ mice. **(C)** *Cln3*^Δex78/WT^ vs. *Cln3*^Δex78/Δex78^ rasters were compared using a permutation sampling method with 1000 trials. Pixels with *p*>0.05 are masked in gray. For regions of significant difference in excitability with *p*<0.05, the difference in fluorescence change (Δ F/FΔex78/WT -Δ F/FΔex78/Δex78) is shown. **(D-E)** In animals expressing *Cln3* in astrocytes (Flex*Cln3* / *Cln3*^Δex78/Δex78^ / *Gfap-*Cre), the DG remained hypoexcitable as compared to the *Cln3*^Δex78/WT^ DG. **(F-G)** Expression of *Cln3* in neurons alone (Flex*Cln3* / *Cln3*^Δex78/Δex78^ / *Syn1-*Cre) was sufficient to reverse hypoexcitability of the DG as compared to *Cln3*^Δex78/WT^ animals. Group sizes (n=slices, N=mice): *Cln3*^Δex78/WT^ n= 23 N=4; *Cln3*^Δex78/Δex78^ n= 30, N=6; Flex*Cln3* / *Cln3*^Δex78/Δex78^ / *Gfap-Cre* n=19 slices, N=4; Flex*Cln3* / *Cln3*^Δex78/Δex78^ / *Syn1-Cre* n=17; N=4.

To evaluate how cell-type specific correction modulates CNS functional defects *in vivo*, we measured network activity through long-term EEG recordings. While restoration of *Cln3* expression in either astrocytes or neurons prevented epileptiform spikes throughout the brain (**Figure 6A**), only rescue in neurons corrected EEG background frequency composition (i.e. frequency band power) and thereby fully normalized network activity **(Figure 6B-C)**. Intriguingly, this correction of physiology occurred despite the presence of substantial storage burden, revealing for the first time that loss of Cln3 protein function in neurons, not storage burden, underlies functional defects in CLN3 disease.

**Figure 6:**
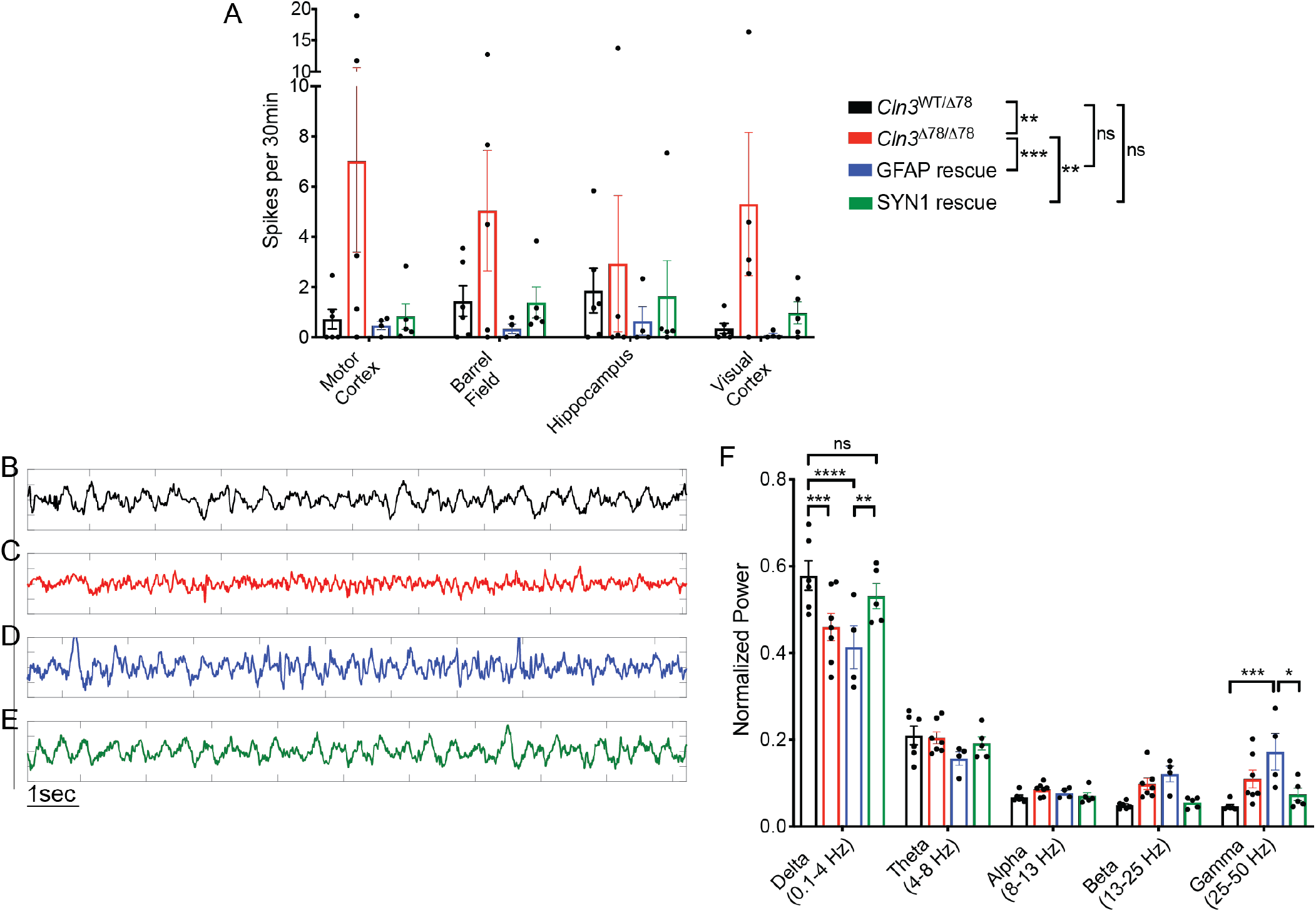
Cln3 rescue in either astrocytes or neurons reduces epileptiform spikes, but only neuronal rescue fully normalizes network dynamics as measured on EEG. **(A)** Correction of either astroctyes (Flex*Cln3* / *Cln3*^Δex78/Δex78^ / *Gfap-*Cre, blue) or neurons (Flex*Cln3* / *Cln3*^Δex78/Δex78^ / *Syn1-*Cre, green) reduced spike burden on EEG in 12 month old mice. **(B-E)** Example EEG traces from *Cln3*^Δex78/Δex78^ (red) and Flex*Cln3* / *Cln3*^Δex78/Δex78^ / *Gfap-*Cre motor cortex (blue) show decreased slow delta activity and increased fast activity as compared to control motor cortex (black). EEG trace from Flex*Cln3* / *Cln3*^Δex78/Δex78^ / *Syn1-*Cre motor cortex (green) shows that rescue of *Cln3* expression in neurons normalizes EEG background frequency composition. **(F)** Power spectral analysis of EEG data recorded in the motor cortex confirms that rescue of *Cln3* expression in astrocytes (blue) does not correct disease-associated loss of delta activity, and actually exacerbates increased fast gamma activity. However, correction in neurons (green) normalizes EEG frequency composition. N=4-5 animals/genotype, after passing a Shapiro-Wilk test of normality, data were analyzed by two-way ANOVA, followed by Sidak’s multiple comparisons test. Significant differences from multiple comparisons testing are shown as \**p*<0.05, \*\**p*<0.01, \*\*\**p*<0.001. Frequency data in panel F in *Cln3*^WT/Δex78^ and *Cln3*^Δex78/Δex78^ also shown in Fig 1D.

## Discussion

Previous cellular-level studies of CLN3 disease have demonstrated disease-associated abnormalities in both neurons and astrocytes^4,7-9,11,26-28^. Prior to this work, it was unknown how these cellular defects translate to the circuit-level dysfunction that underlies neurological symptoms such as dementia and seizures that CLN3 disease patients experience. This work highlights the fact that neurons underlie functional defects in CLN3 disease and must be targeted in future drug development efforts.

A number of novel therapies, including small molecule^5,29-33^, antisense oligonucleotides^34^, and gene replacement ^35,36^ strategies are under development for CLN3 disease. These approaches will not target all cell types equally. For example, studies in non-human primates have demonstrated that intravenous administration of AAV9 in juvenile non-human primates predominately transduces glia^21,37^ while direct administration into the cerebral spinal fluid (CSF) improves delivery to neurons^38^ (34). Our results suggest that maximizing neuronal correction will be essential to develop a treatment that improves functional outcomes in CLN3 disease.

In previous studies, *Cln3*^Δex78/Δex78^ mice have demonstrated abnormal astrocytic-neuronal coupling^7^. Monogenic mouse models of isolated astrocyte dysfunction have epileptiform spikes and seizures on EEG because of pathologic cross-talk between astrocytes and neurons^39^. In the Flex*Cln3* mouse, while rescue of *Cln3* expression in astrocytes did not normalize EEG background, it was sufficient to prevent epileptiform spikes. The reduction in spiking likely reflects improved astrocytic-neuronal interactions when *Cln3* expression is restored in astrocytes. Collectively, our data suggests that diseased astrocytes may modulate network-level dysfunction in CLN3 disease, but diseased neurons are the primary driver of functional deficits.

Using our cell-type specific rescue model, we found that storage accumulation was prevented when *Cln3* was reintroduced to all cell types embryonically, but not when that correction was restricted to astrocytes or neurons alone. Interestingly, prevention of storage accumulation is not required for correcting neuronal network physiology, suggesting that storage material does not underlie functional defects and may be an innocent bystander in the disease process. These studies highlight the importance of using clinically-relevant functional measures, such as *in vivo* electrophysiology studies, to guide treatment development. Histologic measures, such as storage burden, may be poor surrogate biomarkers of therapeutic efficacy in CLN3 disease and possibly other neuronopathic LSDs.

In summary, partial restoration of Cln3 expression by AAV vectors or through Cre-based systems shows that neuronal expression is critical for preventing deficits in CNS functional circuits in CLN3 disease mice. This work also highlights that improvement in storage burden may be a poor readout of therapeutic efficacy in CLN3 disease, a phenomenon that should be considered by the broader LSD community.

## Materials and Methods

For additional information, see Supplemental Methods.

### Study approval

All animal protocols were approved by the Institutional Animal Care and Use Committee at the Children’s Hospital of Philadelphia.

### Statistical Analysis

All statistical analysis of VSDI rasters was completed using a permutation sampling method and the VSDI toolbox in MATLAB (Mathworks) as previously described^12,40^. For summary statistics of regions or groups, GraphPad Prism was also used. In graphs, data are expressed as mean ± SEM. The statistical significance of the observed differences between 2 groups was assessed by two-tailed t-test (for normally distributed data) or Mann-Whitney U test (for non-normally distributed data). For grouped data sets that were not normally distributed a Kruskal-Wallis test followed by a multiple comparisons test was used as detailed in each figure legend. For grouped data sets that were normally distributed, 1-way ANOVA or 2-way ANOVA followed by multiple-comparisons test was completed as detailed in each figure legend. Results were considered significant when *p* < 0.05. Male and female mice were included in all groups.

## Acknowledgments

NINDS R21NS084424 and NICHD HD33532 to BLD, NIH K08NS105865 to RAN, NCL Stiftung Research Award to RAN, Children’s Hospital of Philadelphia Research Institute support to RAN and BLD.

## Conflicts of interests

BLD is a founder of Spark Therapeutics and Spirovant is on the scientific advisory boards of Intellia Therapeutics, Homology Medicines, Prevail Therapeutics, Resilience, Moment Bio, Spirovant, Saliogen and Panorama Medicines. No other authors report conflicts of interest.

## Author contributions

Design of study: RA, LT, AH, OK, RC, EL, EM, CS, BD

Data acquisition and analysis of data: RA, LT, AH, OK, RC, EL, CS

Drafting of manuscript: RA, LT, AH, OK, CS

Critical review of data and manuscript: RA, LT, EM, CS, BD

