## Supplement for "Neuronal genetic rescue normalizes brain network dynamics in a lysosomal storage disorder despite persistent storage accumulation"

**Supplemental Methods:**

*Luciferase assay for promoter validation*

A luciferase assay was established to confirm expression from the 1665bp *Cln3* promoter **(Sup Figure 1)**. Specifically, 1665 bp from the murine Cln3 gene directly upstream of the translational start site (gene ID ENSMUSG00000030720.16; transcript ENSMUST00000032962.10, protein Uniprot Q61124) was obtained from C57BL6/J genomic DNA using PCR with:

Forward primer 5'-GAGGGGGCATTCGGCAAGTGCAGCC -3'

Reverse primer 5'-AACATTGAGTTCGGGTCCCCCAAAGGG-3'.

The p1665 sequence contains an osmotic response element (ORE) [1] at position 646-656 bp and a Coordinated Lysosomal Expression and Regulation (CLEAR) element for TFEB binding at position 1352-1361 bp[2]. The p1665 was initially cloned into pCRBlunt II-TOPO (Invitrogen) and from there was subcloned upstream of firefly luciferase in pGL2-basic vector (Promega) using restriction enzyme digest and standard cloning methods to create p1665luc. As a positive control, the RSV promoter was similarly cloned into pGL2-basic to create RSVluc.

MDCK cells (ATCC) were cultured in DMEM/F12 with 10% FBS. After trypsin harvest, cells were suspended in media at 100,000/ml and transfected with RSVluc or p1665luc plasmid via reverse transfection using Lipofectamine Ltx with Plus reagent (Invitrogen), at a ratio of 400ng plasmid/0.8 ul Lipofectamine Ltx/ 0.4ul Plus reagent/50,000 cells per well, in 24-well culture dishes. The media was changed the next day and three days post transfection cells were harvested into passive lysis buffer (Promega) containing 1x protease inhibitor (200ul lysis buffer per well) and lysates stored at -80°C till luciferase assay. Luciferase activity was determined using Promega luciferase kit reagents. Five microliters of lysate were assayed in duplicate in a luminometer plate reader, with readings collected after injection of 50 ul of substrate. Results are expressed as luminescence units (LU) ± standard deviation of duplicate cultures. To ensure results were within the linear range, lysate from RSVluc-transfected wells (1, 2, 5, 10, 20 ul) was assayed in duplicate on the same plate; all sample readings fell within the linear range.

*AAV9-mediated gene replacement*

Mouse *Cln3* and *eGFP* genes were PCR amplified from cDNA libraries and cloned into the pKAAV.EC.BGHpA backbone plasmid (**Sup Figure 2**). The CAG promoter in the backbone plasmid was substituted by 1665 bp sequence of the endogenous *Cln3* gene.

AAV9 viral vectors expressing *Cln3* or *eGFP* were generated using standard triple transfection methods and purification by CsCl gradient centrifugation [3]. Titers were quantified by silver stain after gel electrophoresis (SDS-PAGE) and qPCR.

Cryo-anesthetized neonatal pups (P0-P1) from *Cln3*$^{\Delta ex78/\Delta ex78}$ X *Cln3*$^{WT/\Delta ex78}$ breeders were ICV injected into the right lateral ventricle with 1E+10 vg AAV9 viral vectors in 2.5 µL according to published methods [4]. In brief, a 10 µL Hamilton microsyringe with a 33-gauge needle was used to penetrate the skull to a 2 mm depth at about 2.5 mm anterior to the lambda suture and 1 mm lateral to the sagittal suture. Injected mice were euthanized, and brain tissue analyzed 1 month or 1 year after injection. We confirmed transduction of both neruons and astrocytes in 1 month old mice after treatment with the AAV9.1665eGFP vector (**Sup Figure 3**). At 12 months of age, gene replacement in *Cln3*$^{\Delta ex78/\Delta ex78}$ lead to a partial reduction of storage accumulation in the cortex (**Sup Figure 4**).

*FlexCln3 mouse generation*

The Flex*Cln3* mouse was generated using previously-published TARGATT technology (Applied StemCell, ASC) [2]. Specifically, a plasmid containing the Flex*Cln3* allele with 1665bp of the endogenous Cln3 promoter was created (**Sup Figure 5**). The targeting construct contained the following features flanked by attB sites: p1665-lox2272(for)-loxP(for)-Cln3 cDNA(rev)-lox2272(rev)-loxP(rev)-BGH polyA. Once transgenic mice are established and bred to cre recombinase driver strains, the lox2277/loxP sequences allow for cre-recombinase mediated flip/excision steps, to place the Cln3 cDNA into forward orientation, triggering expression driven by the p1665 promoter. The targeting construct was used by ASC for microinjection into C57BL6 zygotes for attB-dependent recombination into attP receptor sites in the H11 gene locus.

Successful integration in four founder animals was identified by PCR using a panel of primers **(Sup Figure 6)**. Identification of appropriate founders included validation using 4 different PCR primer pairs:

**PCR 1: PR425N/PR436.** PR425N (forward primer) binds to the unique sequence on H11P3 allele, PR436 binds to partial sequence of attL. P425N/PR436 pair is to detect site-specific integration at 5' integration site. Expected PCR fragment is 217bp for 2, 3 insertion or 147bp for 1, 2 or 1, 3 insertion.

**PCR 2: PR425N/1665-R.** PR425N binds to the unique sequence on H11P3 allele, while 1665-R (reverse primer) recognizes the sequence within 1665 promoter region. A combination of PR425N and 1665-R can be used to amplify 5' novel junction upon the successful integration of transgene. Anticipated DNA amplification fragment is 495 for 2, 3 insertion, or 425bp for 1, 2 or 1, 3 insertion.

**PCR 3: BGH-pA-F/PR387**: BGH-pA-F binds to BGH polyA sequence, while PR387 (reverse primer) binds to H11 genomic sequence downstream of 3' junction. The PCR amplification using BGH-pA-F/PR387 pair produces a 393bp (1, 3 or 2, 3 insertion) or 463bp (1, 2 insertion) fragment with part of BGH-polyA and partial H11 genomic sequence.

**PCR 4: PR522/SH178.** PR522 binds to partial sequence of attR. PR178 binds to a region in H11P3 locus downstream of the integration site. PCR amplification using PR522/PR178 pair is to detect site-specific integration at 3' integration site. Expected PCR fragment is 417bp for 2, 3 or 1, 3 insertion, or 487bp for 1, 2 insertion.

*Generation of cell-type specific rescue models*

Founder Flex*Cln3* mice were confirmed by PCR and one founder was selected to maintain the transgenic line **(Sup. Figure 6**). The transgenic line, mice harboring the Flex*Cln3* allele, as well as the Cre recombinase driver strains used in this study, were crossed to the well-established *Cln3*$^{\Delta ex78}$ mouse model[5] (Jackson Labs, B6.129(Cg)-*Cln3*$^{tm1.1Mem}$/J, Strain #017895). This model harbors the most common human CLN3 disease mutation, and manifests histopathologic features of CLN3 disease including progressive storage accumulation. For these experiments, 3 Flex*Cln3* lines were generated: 1) To express the Flex*Cln3* in all cell types (ubiquitous rescue), Flex*Cln3* / *Cln3*$^{\Delta ex78/\Delta ex78}$ were crossed to mice expressing Cre recombinase under the control of the E2a (B6.FVB-Tg(EIIa-cre)C5379Lmgd/J, Strain # 003724) or *Zp3* promoter [6] (*Zp3-Cre,* Jackson Labs, B6.Cg-Tg(Zp3-cre)1Gwh/J, Strain #06888) for recombination at germline level 2) To selectively express the wild-type Flex*Cln3* transgene in neurons, Flex*Cln3* / *Cln3*$^{\Delta ex78/\Delta ex78}$ mice were crossed to previously described mice expressing Cre recombinase under control of the Synapsin-1 promoter[7] (*Syn1-Cre*, Jackson Labs, B6.Cg-Tg(Syn1-cre)671Jxm/J, Strain #003966). 3) To express wild-type Flex*Cln3* in astrocytes, Flex*Cln3* / *Cln3*$^{\Delta ex78/\Delta ex78}$ mice were crossed to previously described mice expressing Cre recombinase under control of the glial fibrillary acidic protein promoter[8] (*Gfap-Cre*, Jackson Labs, B6.Cg-Tg(Gfap-cre)77.6Mvs/2J, Strain #024098).

*Verification of Cre-mediated transgene flipping in genomic DNA from target tissues*

For each line, tissue-specific Cre-mediated recombination of the Flex*Cln3* allele was confirmed through PCR of genomic DNA extracted from liver, spleen and brain **(Fig 2, 3 and Sup. Figure 7)**. For initial confirmation of Flex*Cln3* allele recombination and flipping in target tissues, genomic DNA was extracted from Flex*Cln3* / *Cln3*$^{\Delta ex78/\Delta ex78}$ mice crossed to the Cre-driver lines above.

For confirmation of orientation of the Flex*Cln3* allele via PCR of genomic DNA from target tissues the following primers were used:
Reverse orientation of the Flex*Cln3* transgene:
Forward primer: 5'-TGGGGCTCTGAGTCGGTCTC-3'
Reverse primer: 5'-TGGGAGTGGCACCTTCCAGG-3'
Product size 380 bp.

Forward orientation of the Flex*Cln3* transgene
Forward primer: 5'-GGAGACCAGTGACAAGCACCG-3'
Reverse primer: 5'-TGGGAGTGGCACCTTCCAGG-3'
Product size 345 bp.

*Expression of FlexCln3 transcripts*
*Cln3* expression was quantified from the whole hippocampus of 4-month-old mice. Samples were flash-frozen in 200 µL of TRIzol reagent (Invitrogen, ThermoFisher Scientific) per sample. RNA was isolated in 1mL TRIzol followed by DNase treatment with TURBO DNA-free kit (Invitrogen, ThermoFisher Scientific), according to manufacturer's protocols. RNA concentration was calculated and purity tested for 260/280 ratio greater than 1.9 by NanoDrop 2000 (ThermoFisher Scientific). cDNA was synthetized from 2 µg RNA by reverse transcription using Superscript III or High capacity cDNA reverse transcription kit (Life Technologies, Carlsbad, CA) with RT random primers according to the manufacturer's recommendations. Quantitative PCR reactions were performed on a CFX384 Real Time System (Bio-Rad, Hercules, CA). A custom PrimeTime primer/probe assay (IDT-integrated DNA Technologies (Coralridge, IA) was used for amplification of forward-oriented FLEX*Cln3* transcripts off the transgenic allele:

Forward primer: 5'-GCTCGAGCATGCATCTAGTATAA-3'
Reverse primer: 5'-AAGGCACAGTCGAGGTCTA-3'
Probe: 56-FAM/AGGTTCGAA/ZEN/TTCGATATCGCGGCC/3IABkFQ

A commercially available primer/probe assay for murine *Cln3* (IDT assay Mm.PT.58.9756823) recognizing wildtype (exon 7/8-containing) transcripts was used to amplify *Cln3* transcripts from control mice (*Cln3*$^{\Delta ex78/WT}$or *Cln3*$^{WT/WT}$) and from the FLEX*Cln3* locus of ubiquitously rescued mice. *ACTB* (control mix-VIC, Applied Biosystems, #4351315) was used as a reference gene to normalize Cln3 expression levels by the delta delta cycle threshold (ddCt) method.

*Western Blot for subunit C of the mitochondrial ATP synthase (SCMAS)*

Mice were euthanized under anesthesia, the brain removed and regions isolated by dissection in ice-cold PBS under a dissecting microscope, and snap frozen in liquid nitrogen. Samples were stored at -80°C until processing. Tissues were processed by homogenization in Eppendorf microcentrifuge tubes in ice-cold lysis buffer (50 mM Tris pH 7.5, 150 mM NaCl, 1% NP40, 0.1% SDS, 0.5% DOC, with 1x Roche protease inhibitors). After 20 min incubation on ice, lysates were centrifuged at 1600xg for 5 min at 4 °C to pellet debris, and lysates transferred to fresh tubes. Protein concentrations were determined (BioRad DC assay) and concentrations equilibrated to 2 mg/ml, and 450 ul of each lysate was centrifuged at 20,817xg for 30 min at 4 °C. Supernatants (detergent-soluble fraction) were transferred to fresh tubes. Pellets (detergent-insoluble fraction) were dispersed in 500 ul lysis buffer and the centrifugation step repeated. Washed pellets were resuspended in 90 ul lysis buffer supplemented with 1% SDS and 1x NuPage and sonicated for 3 s on ice. NuPage loading buffer was added to the detergent-soluble samples.

All samples were heated at 70 °C for 10 min and 16 ul/lane loaded onto precast NuPage 4-12% Bis-tris mini-gels, with Spectra Multicolor Broad Range protein ladder (Thermofisher), run in MES buffer, and transferred overnight in a Mini Trans-Blot Cell (BioRad) in Tris-glycine transfer buffer containing 10 % methanol and 0.01% SDS at 4 °C with constant 30 V onto PVDF membranes. Membranes were blocked by 1 h incubation in 5% dried milk powder in PBS with 0.05% tween (PBST), then cut into strips for overnight incubation at 4 °C with primary antibody diluted in 2.5% dried milk in PBST. Antibodies to the following antigens were used for blotting: SCMAS at 1/3000 (polyclonal rabbit antisera from Elizabeth Neufeld); p62 at 1/4000 (Abnova #H00008878-MO1 mouse IgG clone 2C11, 0.5 mg/ml stock); beta-actin at 1/20,000 (Sigma mouse IgG clone AC-15 ascites). After primary antibody incubation, blots were washed in PBST and incubated with HRP-conjugated secondary goat-anti-rabbit IgG or goat-anti-mouse IgG (Jackson ImmunoResearch) for 1 h at RT. Blots washed and developed with ECL prime (Amersham) and chemiluminescent images captured and analyzed using a Versadoc image capture system with Quantity One software (Bio-Rad). Beta-actin was used as a loading control, and band densities for SCMAS and p62 were normalized to beta-actin in each lane. Graphs show the relative mean band intensity ± standard deviation between groups.

*Histology*

Immunohistochemical analysis of fixed brains was carried out as previously described[9]. Mice were anesthetized with isoflurane and transcardially perfused with saline and 4% paraformaldehyde (PFA) in 0.1 M phosphate buffer (MilliporeSigma), pH 7.4. The brains were then postfixed in 4%PFA at 4°C overnight, cryoprotected with 30% sucrose buffer and embedded in OCT. Coronal sections (20 μm) were cut using a cryostat (Microm HM 500) and stored at –80°C until staining.

For mitochondrial ATP synthase subunit C (SCMAS) staining, sections were washed with PBS, permeabilized with 0.1% Triton X-100 in PBS for 7 minutes, washed again with PBS 3 times, and blocked in 3% bovine serum albumin, 5% goat serum, 0.2% Triton X-100 in PBS for 1 hour, all at room temperature. Sections were incubated with a primary antibody against subunit C of the mitochondrial ATP synthase (SCMAS) (Abcam 181243, 1:500).

For GFAP staining, sections were washed with PBS, permeabilized with 0.1% Triton X-100 in PBS for 7 minutes, incubated in 3% H2O2/10% methanol in PBS for 5 minutes, washed again with PBS 3 times, and blocked in 10% normal goat serum with 0.1% Triton X-100 in PBS for 1 hour, all at room temperature. Sections were incubated with a primary antibody against GFAP (Dako Z0334, 1:1000) at 4°C overnight.

For NeuN staining, sections were washed with PBS, permeabilized with 0.1% Triton X-100 in PBS for 7 minutes, incubated in 3% H2O2/10% methanol in PBS for 5 minutes, washed again with PBS 3 times, and blocked in 10% normal goat serum in PBS, all at room temperature. Sections were incubated with an antibody against NeuN (MilliporeSigma MAB377, 1:100) at 4°C overnight.

To detect the primary antibody using DAB staining, sections were incubated with a biotinylated secondary antibody (Jackson ImmunoResearch, 1:500) at room temperature for 30 minutes, followed by PBS washes and incubation in avidin-biotin complex (Vectastain, Vector Laboratories) at room temperature for 1 hour. Slices were incubated in DAB solution for visualization.

To detect the primary antibody using immunofluorescence, sections were incubated with a FITC-conjugated secondary antibody (Jackson ImmunoResearch, 1:50) at room temperature for 30 minutes, followed by PBS washes. For immunofluorescence, all auto-fluorescence was quenched by incubation for 30 seconds in TrueBlack (Biotium, 50ul of 20X stock in 1ml of 70% ethanol).

All images were acquired on a Keyence BZ-X800 or a Leica DM600B microscope and analyzed in a blinded fashion. At least 4 animals per genotype were analyzed. For quantification of SCMAS staining, Keyence BZ-X800 analysis software was used to calculate the percent area of fluorescence. In the CA3 region, one 10X image of the entire hippocampal CA3 region was captured per animal, while in the motor cortex at least 4, 20X images/animal were acquired. For GFAP staining, at least 4, 20X images / animal were acquired for both the motor cortex and CA3 regions. Images were converted to 8-bit grayscale images and the integrated density of the image was calculated in ImageJ, using rolling ball radius background subtraction.

*Voltage Sensitive Dye Imaging*

*In vitro* voltage sensitive dye imaging (VSDI) of hippocampal slices was completed as previously described[9]. 400μm hippocampal-entorhinal cortical (HEC) slices, which preserve the hippocampal trisynaptic circuit [10-14], were prepared as per our standard protocols[15]. High-sucrose cutting solution contained in mM, 192 sucrose, 2.5 KCl, 1.25 $NaH_2PO_4$, 26 $NaHCO_3$, 12.2 glucose, 3 sodium pyruvate, 5 sodium ascorbate, 2 thiourea, 10 $MgSO_4$, 0.5 $CaCl_2$. After cutting, slices recovered for 45 min at 37°C and 45 min at room temperature prior to recordings in an artificial cerebrospinal fluid (ACSF) containing, in mM, 115 NaCl, 2.5 KCl, 1.4 $NaH_2PO_4$, 24 $NaHCO_3$, 12.5 glucose, 3 sodium pyruvate, 5 sodium ascorbate, 2 thiourea, 1 $MgSO_4$, 2.5 $CaCl_2$. For recording, slices remained in a standard ACSF solution containing, in mM, 128 NaCl, 2.5 KCl, 1.4 $NaH_2PO_4$, 26.2 $NaHCO_3$, 12.2 glucose, 1 $MgSO_4$, 2.5 $CaCl_2$.

The VSD di-3-ANEPPDHQ (Invitrogen) was solubilized in 95% ethanol (0.020 mg/uL) and stored at -20°C. Slices were stained with the di-3-ANEPPDHQ (0.1 mg/mL, in ACSF) for 14-16 minutes and transferred to a humidified interface chamber (BSC2, Scientific Systems Design), for recording. Excitation light was provided by 7 high-power green LEDs (Luxeon Rebel LXML-PM01-0100, Philips) coupled to a 535 ± 25 nm filter and 565 nm dichroic mirror. A 610 nm longpass filter further isolated the emitted fluorescence. Fluorescence was recorded at 1000 frames per second with a fast video camera with 80 x 80 pixel resolution (NeuroCCD, Redshirt Imaging, Decatur, GA).

For experiments, four 0.1ms pulses at 10Hz were delivered via a bipolar tungsten electrode (model ME12206, World Precision Instruments) positioned to stimulate the perforant pathway.
Stimulus strength was set at the current required to produce a saturating field potential response in the granule cell layer of the dentate gyrus using a glass electrode pulled to a 2-8 MΩ tip, and filled with ACSF. All recordings were 1.5 s long, with a 10 s delay between recordings to allow fluorescence to recover from photobleaching. Thirteen recordings of evoked activity were interleaved with 13 VSDI runs

where no stimulus was delivered. Runs without delivered stimuli were used for offline subtraction to correct for any baseline drift over the course of a recording.

Post-collection VSDI analysis was completed using the previously published VSDI toolbox [15], which allows for robust statistical analysis across both spatial and temporal dimensions. The algorithm creates unbiased regions of interest over the entire dentate gyrus, and the fluorescence response in each ROI for each image is calculated. As previously published [15], in order to compare between and average slices linear interpolation was used in order to "stretch" rasters to be the same size. For all studies, a final stretched DG raster size was 22 ROIs, including 11 covering the internal and external blades each.

A two-dimensional raster plot showing the location, time and fluorescence change for each slice was generated. In addition to individual rasters for each slice, an average raster plot per condition was generated. The raster plots from different genotypes were compared using a permutation sampling method with n=1000 permutations. A heatmap showing regions of the network with statistically significant ($p<0.05$) differences was created. Group sizes for VSD studies were as follows: *Cln3*$^{\Delta ex78/WT}$ 23 slices from 4 animals; *Cln3*$^{\Delta ex78/\Delta ex78}$ 30 slices from 6 animals; Flex*Cln3 / Cln3*$^{\Delta ex78/\Delta ex78}$ / *Gfap-Cre* 19 slices from 4 animals; Flex*Cln3* / *Cln3*$^{\Delta ex78/\Delta ex78}$ / *Syn1-Cre* 17 slices from 4 animals.

*EEG acquisition and analysis*

EEG recording electrodes were constructed and implanted as previously described [9]. Specifically, recording headcaps were created by attaching 1 ground, 1 reference, and 6 cortical leads lead (0.004 inches, formvar-coated silver, California Fine Wire), and 2 hippocampal leads (0.005 inches bare, 0.008 inches coated, stainless steel wire, AM-Systems) to a microconnector (Omnetics). For implantation, mice were premedicated with intramuscular ketamine/xylazine followed by inhaled isoflurane anesthesia. The following stereotaxic coordinates (measurements relative to bregma) were used to implant electrodes: bilateral motor cortices: 0.5 mm rostral, 1 mm lateral, and 0.6 mm deep; bilateral barrel field cortices: 0.7 mm caudal, 3 mm lateral, and 0.6 mm deep; bilateral visual cortex: 3.5 mm caudal, 2 mm lateral, and 0.6 mm deep; and bilateral hippocampus (CA1 region): 2.2 mm caudal, 2 mm lateral, and 1.7 mm deep. The cerebellar reference lead was implanted posterior to lambda. The ground wire was wrapped tightly around a 1/8-inch self-tap screw (Precision Screws and Parts), which was inserted into the skull rostral to the motor cortex leads. To aid in securing the recording cap, a second screw was inserted caudal to lambda. The recording electrodes were secured with dental cement. The animal was allowed to recover for at least 18 hours before recording. Each mouse was connected to a RHD2000 low-noise amplifier chip (Intan Technologies), and connected to a data acquisition board by a lightweight cable. Electrophysiologic data was acquired at 2 kHz using the Intan RHD2000 Recording system software (http://intantech.com/downloads.html). 12-15 month old animals from each genotype group (n=4-5 animals/group) were recorded for at least 24 hours.

All EEG analysis was completed in MATLAB as previously described[9].

*EEG Analysis*

EEG was analyzed as previously described [9]. Steps included:

1) Detection of poor-quality recording channels: An automated artifact detector was used on each recording channel for each 30-minute recording period before analysis. Detection of a noisy or poorly recording channel was completed by calculating the average of the root mean squared amplitude and skew of the voltage for each second of the recording. Channels with a root mean square amplitude of less than 30 μV or more than 200 μV or a skew greater than 0.4 were excluded from further analysis.

2) Spike detection: A spike detector algorithm designed to detect voltage deflections that were greater than 5 standard deviations above the mean with a full width at half maximum amplitude of 5–200 ms was used to detect spikes. Spikes were eliminated if they occurred within 10-seconds of a deflection found to be an artifact (i.e., it had a half width outside the 5- to 200-ms range). Multi-channel spikes were defined as a spike occurring in at least 2 of 8 EEG channels.

3) Power analysis: To analyze mutation-induced discrepancy in background EEG frequency composition, recordings were divided into 5-second epochs for power analysis. Epochs containing large amplitude artifacts, most likely due to movement (defined as a peak *Z*-score of the root mean squared voltage amplitude greater than 3), were excluded from frequency analysis. Fast Fourier transform analysis was then completed on artifact-free epochs. Quantification of the power in each of the major EEG frequency bands (delta 0.1–4.0 Hz, theta 4–8 Hz, alpha 8–13 Hz, beta 13–25 Hz, and gamma 25–50 Hz) was completed. Power in each band across all epochs was averaged and power in each band was normalized to total power.

**Supplemental Figure 1: Luciferase assay confirmation of expression from the 1665bp *Cln3* promoter.** Luminescence was measured from plasmids expressing luciferase under control of the RSV promoter (red) and the *Cln3* 1665bp promoter (blue), as compared to a eGFP control.

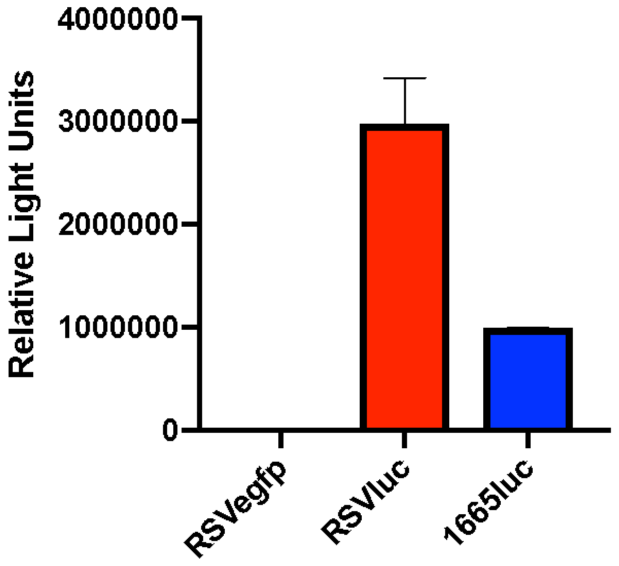

**Supplementary Figure 2: AAV9-CLN3 and AAV9-GFP constructs**

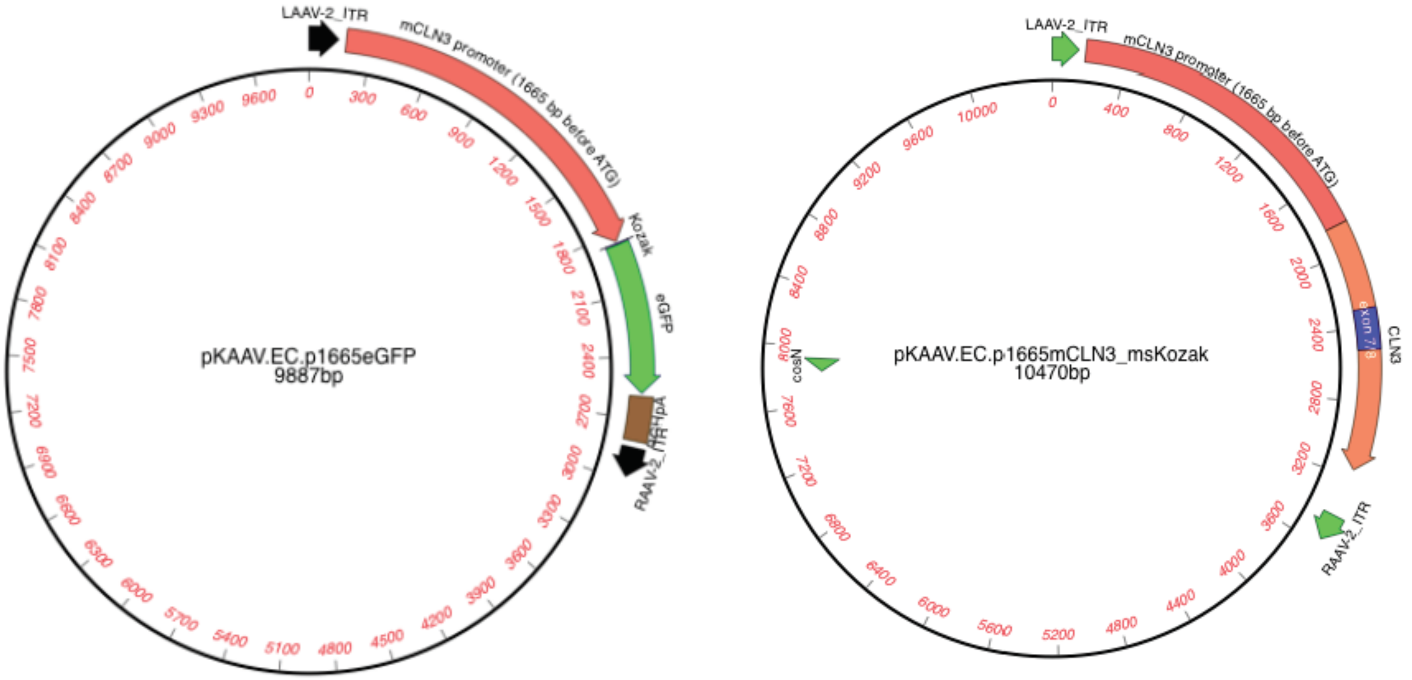

**Supplemental Figure 3: p0 ICV injection of AAV9.1665GFP transduces both astrocytes and neurons.** An AAV9 construct expressing GFP under the control of the 1665bp *Cln3* promoter was given via ICV injection into p0 wildtype pups. In 1-month-old mouse **(A)** cortex and **(B)** hippocampus, neurons, labeled with NeuN, were also GFP positive (examples shown by arrows). In addition, there were NeuN-negative, GFP positive astrocytes detected (examples shown by arrowheads). Sections from AAV.1665CLN3-treated animals that do not express GFP are shown as a negative control.

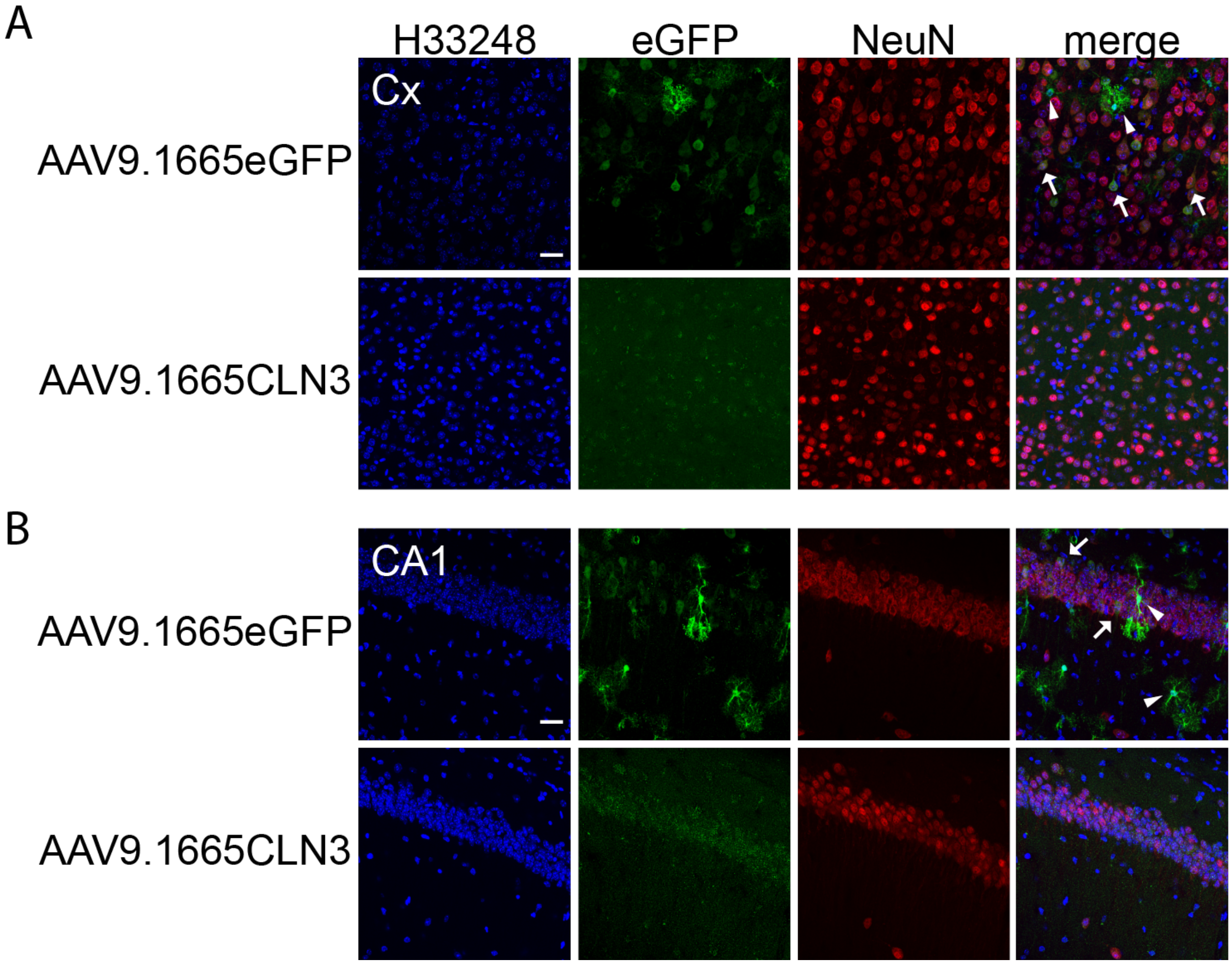

**Supplemental Figure 4: p0 ICV AAV9-CLN3 injection reduces storage accumulation in the motor cortex of 12-month-old *Cln3*$^{\Delta ex78/\Delta ex78}$ mice.** Staining for subunit C of the mitochondrial ATP synthase reveals reduced storage accumulation in AAV9.CLN3-treated *Cln3*$^{\Delta 78/\Delta 78}$ mice as compared to untreated controls.

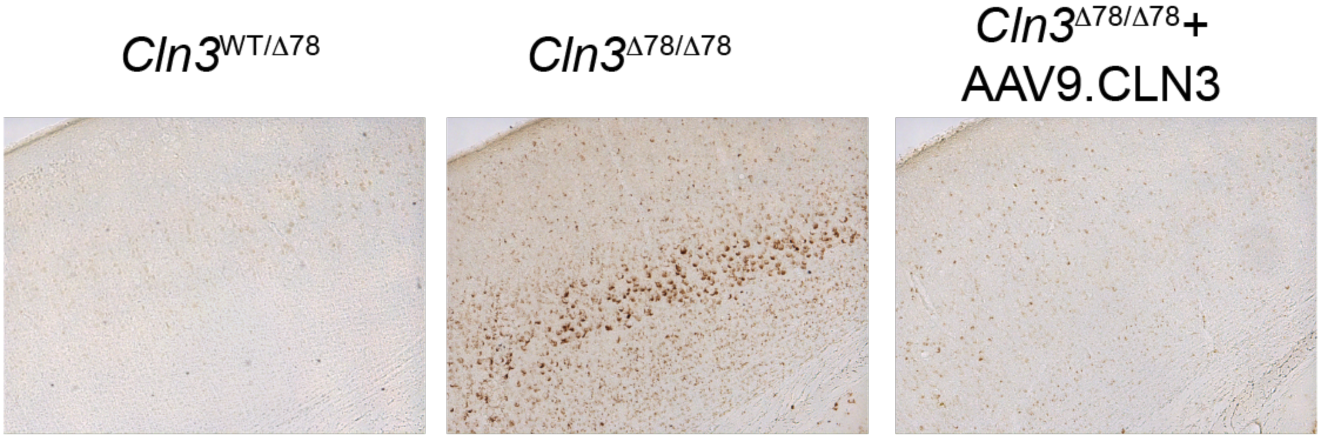

**Supplemental Figure 5:** Map of targeting vectors, pBT378-1665-FLEX-CLN3

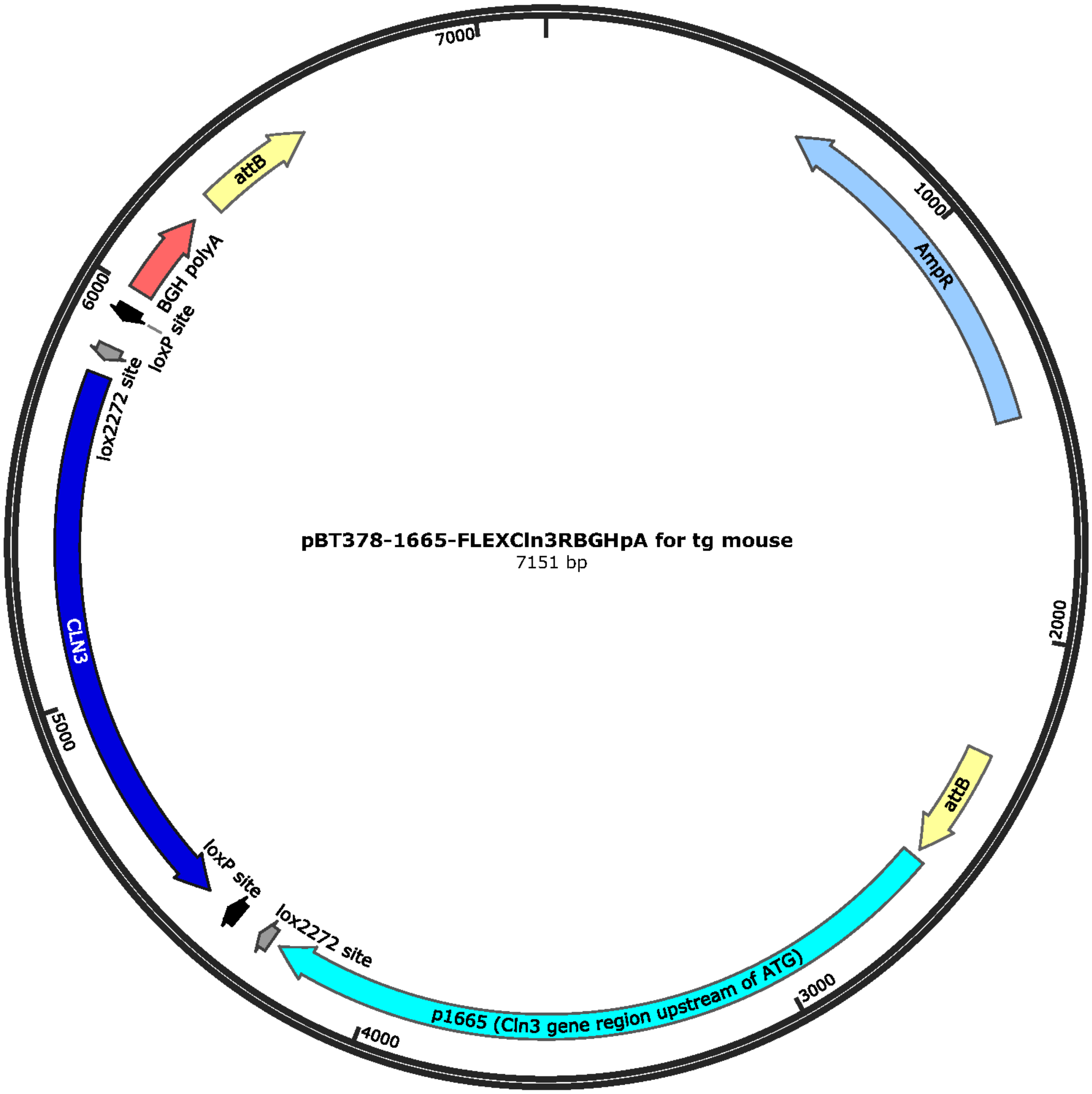

**Supplemental Figure 6:** Confirmation of Flex*Cln3*transgenic founders and first cross to *Cln3*$^{\Delta78/\Delta78}$ mice. **(A)** Schematic shows targeted H11P3 allele without and with transgene insertion. Insertion occurs at a pair of attP sites; insertion at attP1-2 is shown. PCR primers are indicated by arrows (forward in purple and reverse in green), and PCR reactions designated with blue lines. **(B)** Primer sequences and expected product sizes for PCR reactions. **(C)** PCR results from tail-snip DNA confirm transgene insertion for founder male 547#5 (at attP2-3) and for founder female 547#11 (at attP1-3). **(D)** PCR results from tail-snip DNA from litters of first cross of founders to Δex7/8. PCR4 indicates 5 of 29 pups have inherited the FLEXCln3R transgenic allele (417 bp band). Absence of transgene positive progeny from founder 547#5 indicates lack of germline transmission in this founder.

A

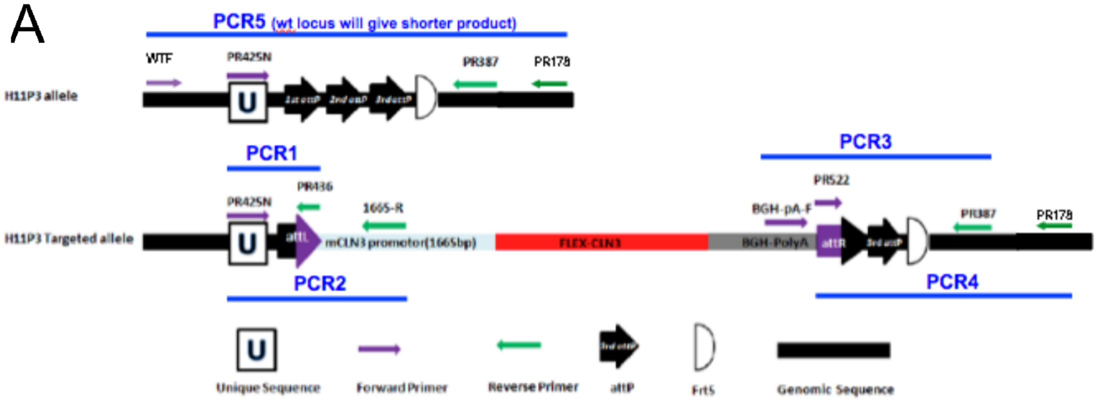


B

| | insertion site | expected PCR product size |
|---|---|---|
| PCR1 | attP 1-2 | 147 bp |
| | attP 1-3 | 147 bp |
| | attP 2-3 | 217 bp |
| PCR2 | attP 1-2 | 425 bp |
| | attP 1-3 | 425 bp |
| | attP 2-3 | 495 bp |
| PCR3 | attP 1-2 | 463 bp |
| | attP 1-3 | 393 bp |
| | attP 2-3 | 393 bp |
| PCR4 | attP 1-2 | 487 bp |
| | attP 1-3 | 417 bp |
| | attP 2-3 | 417 bp |
| PCR5 | no insertion | 660 bp |
| | WT allele | 323 bp |

Primers for genotyping FLEXCln3 founders and progeny
PCRPR425N: 5'-GGTGATAGGTGGCAAGTGGTATTCCGTAAG -3'
PR436: 5'-ATCAACTACCGCCACCTCGAC -3'
1665-R: 5'-CAAACCCTAGCTTCCTCTCACTGAAG-3'
BGH-polyA-F: 5'-GCATCGCATTGTCTGAGTAGGTGTCA-3'
PR387: 5'-GTGGGACTGCTTTTTCCAGA-3'
PR522: 5'-CGATGTAGGTCACGGTCTCG -3'
PR178: 5'-TTCCCTTTCTGCTTCATCTTGC -3'
WTF: 5'-TGGAGGAGGACAAACTGGTCAC-3'

C

| lane | mouse |
|---|---|
| 1 | 547#5 |
| 2 | 547#11 |
| 3 | WT mouse |
| 4 | no template control |

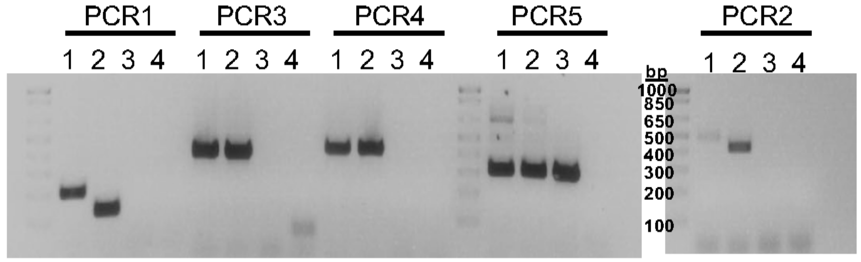


D

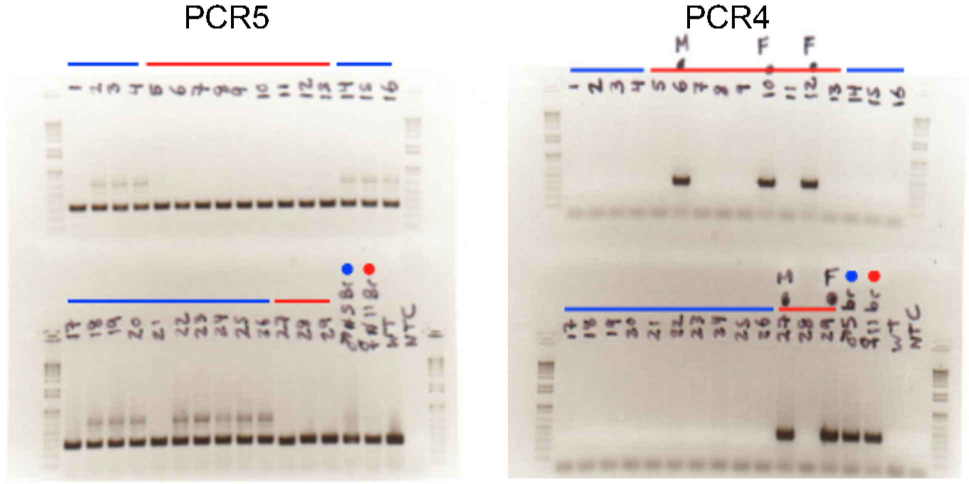


founder parent is 547#5 male
founder parent is 547#11 female

**Supplemental Figure 7: Cre-mediated recombination of Flex*Cln3* allele in target tissues.**
**(TOP)** Full uncut gels showing appropriately rearranged Flex*Cln3* transgene in genomic DNA extracted from hippocampus of ZP3-Cre+, GFAP-Cre+, and Syn1-Cre+ FlexCln3 animals. Lanes shown in Figures 2b and 3a are outlined and labeled. **(BOTTOM)** To confirm that the Flex*Cln3* allele rearranges appropriately in response to a number of Cre drivers, genomic DNA was extracted from the liver, spleen and brain of animals crossed to the drivers listed below. The Flex*Cln3* allele was detected to be in the forward orientation, indicating recombination occurred, in all tissue types in animals crossed to ubiquitous or endothelial Cre lines (lanes 1 and 2). When the Flex*Cln*3 animal was crossed to the neuronal Cre driver, Syn1, (lane3) recombination only occurred in the brain. Lane 4 demonstrates that in the absence of Cre, the allele remains in the reverse orientation.

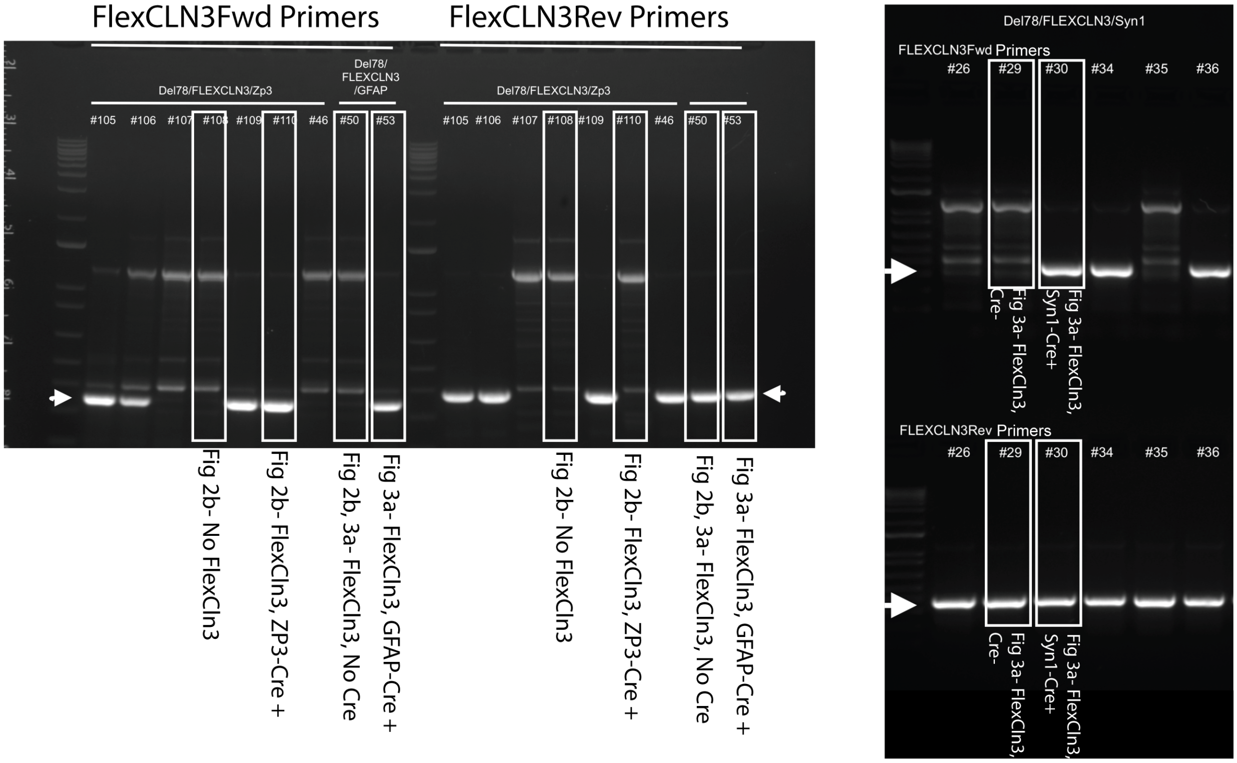


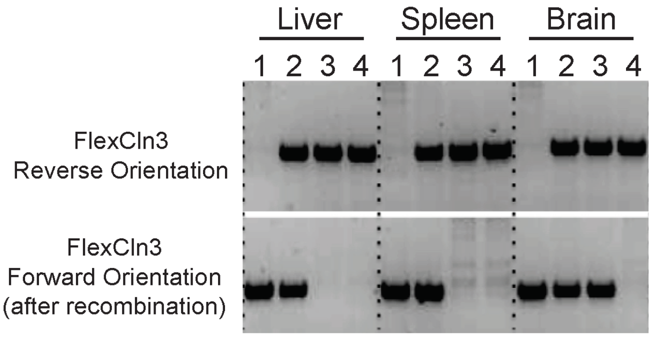


1: Δ78/FlexCln3/E2a-Cre (ubiquitous rescue)
2: Δ78/FlexCln3/Tie2-Cre (endothelial cell rescue)
3: Δ78/FlexCln3/Syn1-Cre (neuronal rescue)
4: Δ78/FlexCln3 (no rescue)

**Supplemental Figure 8: Storage accumulation in Flex*Cln3* mouse.** A. Western blot of NP40 detergent insoluble SCMAS and p62 protein in regional lysates from brain (N=4 per group).

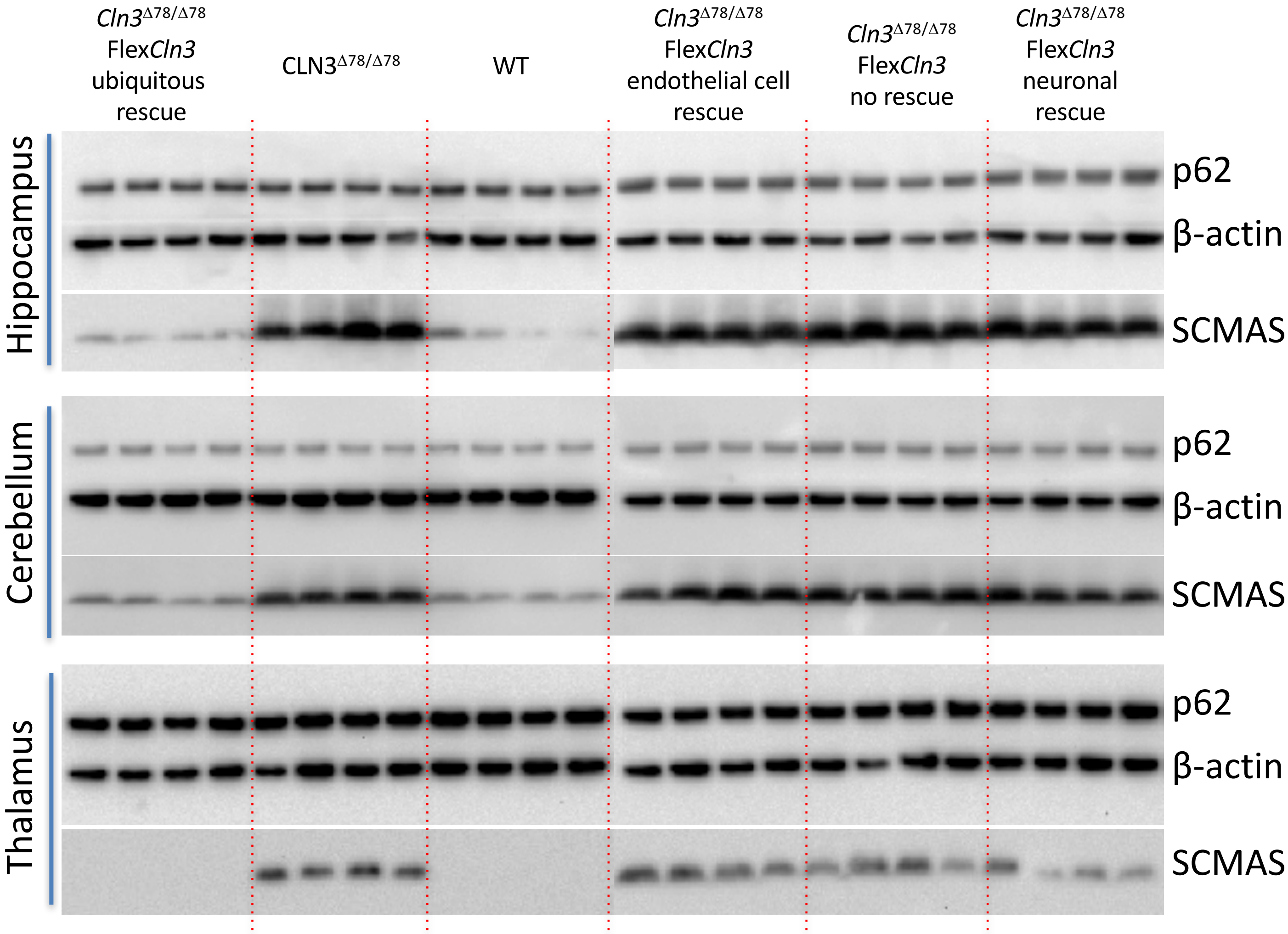

**Supplemental Figure 9: Rescuing expression of *Cln3* in either astrocytes or neurons alone does not prevent storage accumulation in the hippocampus at 6 months of age. (A)** 10X image of the CA3 region of six-month-old *Cln3*$^{WT/WT}$ hippocampus demonstrates little storage of subunit C of the mitochondrial ATP synthase (SCMAS) (green), as compared to **(B)** *Cln3*$^{\Delta ex78/\Delta ex78}$ hippocampus. **(C)** In Flex*Cln3* / *Cln3*$^{\Delta ex78/\Delta ex78}$ / *Gfap*-Cre animals, rescue of *Cln3* expression in astrocytes alone does not prevent SCMAS accumulation. **(D)** Expression of Flex*Cln3* in neurons alone (Flex*Cln3* / *Cln3*$^{\Delta ex78/\Delta ex78}$ / *Syn1*-Cre animals) also does not prevent storage accumulation.

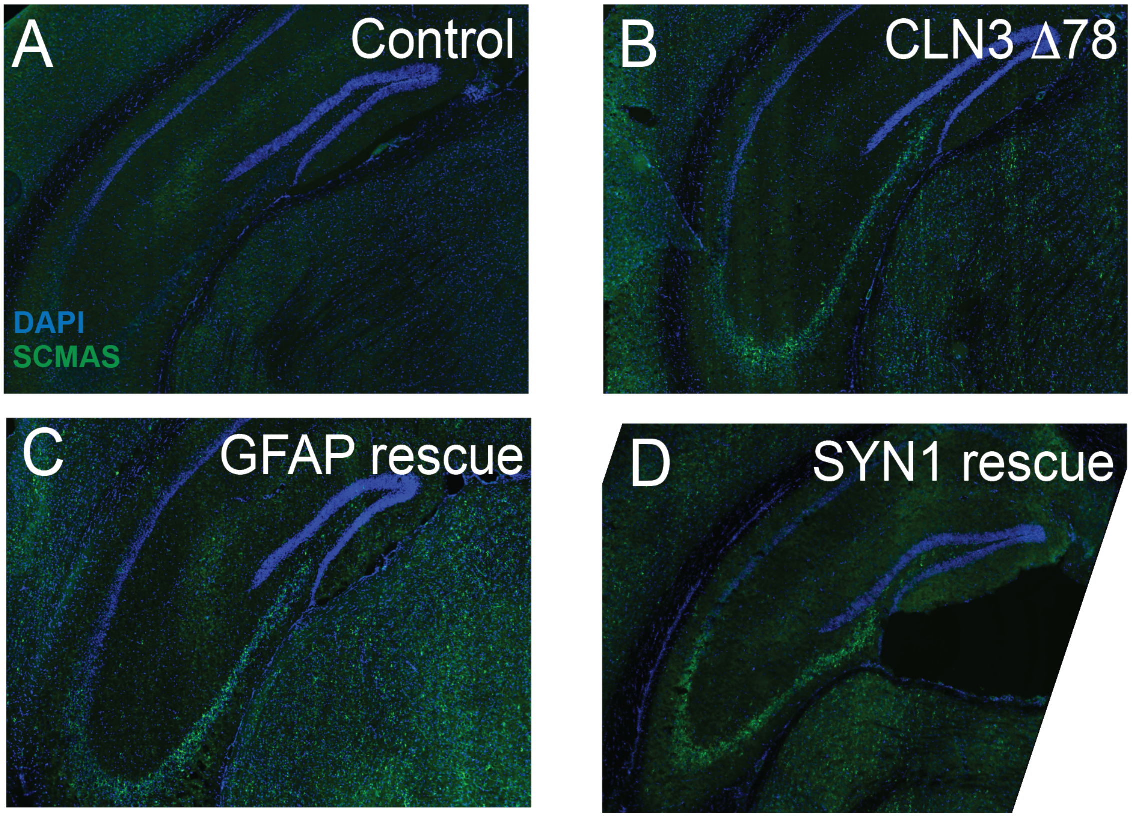

**Supplementary Figure 10: Rescuing expression of *Cln3* in either astrocytes or neurons alone only partially improves astrocytosis in 12-month-old Flex*Cln3* mice. (A-B)** 20X image of the CA3 region of *Cln3*$^{\Delta ex78/\Delta ex78}$ hippocampus demonstrates increased astrocytosis, as measured by GFAP staining as compared to *Cln3*$^{WT/WT}$ hippocampus. **(C-E)** Flex*Cln3* expression in astrocytes alone did not prevent astrocytosis in the CA3 region of the hippocampus. However, *Cln3* expression in neurons alone partially improves astrocytosis. **(F-G)** In the motor cortex, *Cln3*$^{\Delta ex78/\Delta ex78}$ mice demonstrate increased astrocytosis, as compared to *Cln3*$^{WT/WT}$ controls. **(H-J)** Flex*Cln3* expression in either astroctyes of neurons alone partially normalizes astrocytosis in the motor cortex. For both regions, n=4 animals, 6 images / animal, two-way ANOVA by animal and genotype, p<0.0001 for genotype, followed by Tukey's multiple comparisons test. For all panels: *$p$<0.05, **$p$<0.01, ***$p$<0.001, ****$p$<0.0001.

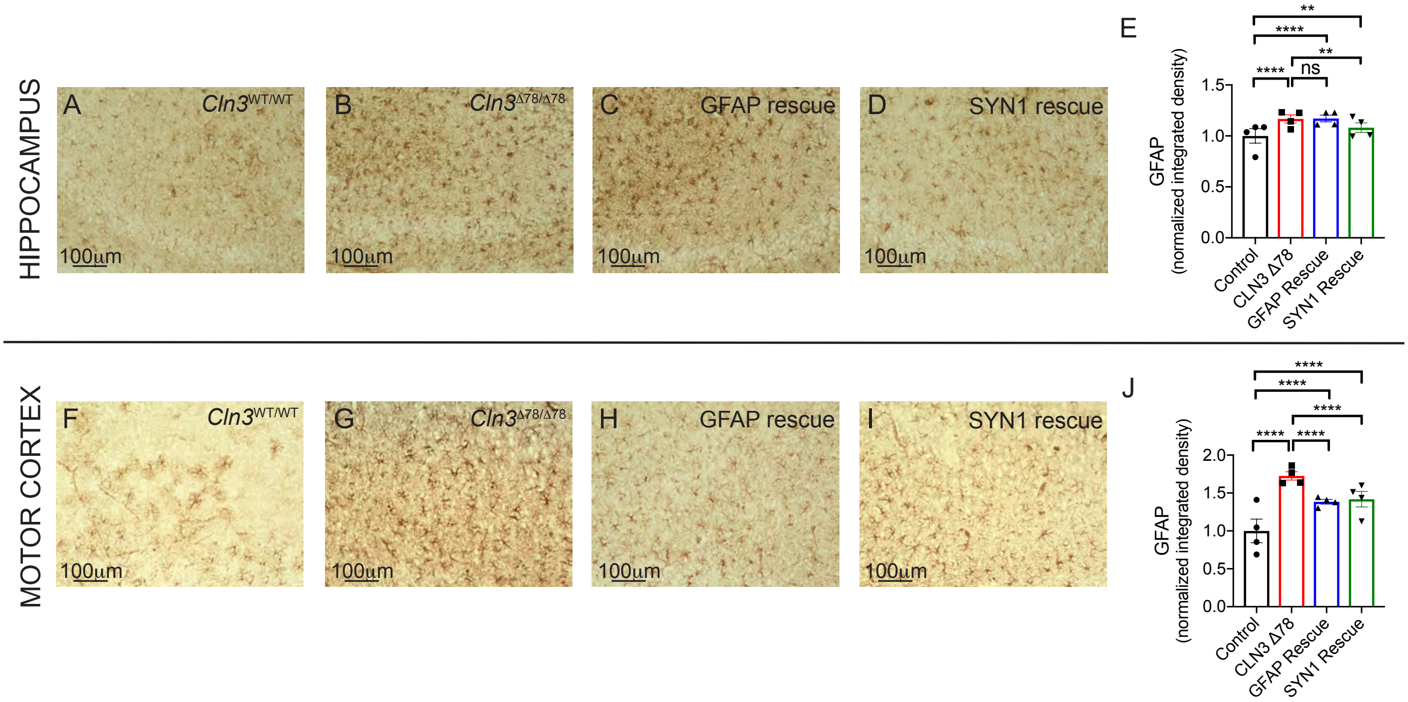
